# A single-cell atlas of transcriptome changes in the intestinal epithelium at the suckling-to-weaning transition

**DOI:** 10.1101/2024.12.02.626361

**Authors:** Tania Malonga, Christelle Knudsen, Emeline Lhuillier, Patrick Aymard, Elisabeth Jones, Corinne Lencina, Manon Despeyroux, Elodie Riant, Cédric Cabau, Alyssa Ivy, Crystal L. Loving, Nathalie Vialaneix, Martin Beaumont

**Author notes:** Correspondence, Address: 24 Chemin de Borde Rouge, 31320 Castanet Tolosan, France. Contributed equally (Nathalie Vialaneix, Martin Beaumont).

## Abstract

The suckling-to-weaning dietary transition is a key step in mammalian intestinal development. However, the relative contributions of genetically wired and nutritional factors in this maturation process remain to be elucidated. Moreover, the cellular diversity of the intestinal epithelium has been overlooked in this context. The aim of our study was to identify the transcriptome changes induced in each cell type of the intestinal epithelium at the onset of solid food ingestion. We compared the single-cell transcriptome of epithelial cells isolated from the caecum of age-matched littermate suckling rabbits ingesting or not solid food. Our dataset provides the first single-cell atlas of the rabbit intestinal epithelium and highlights the interest of the rabbit as a model for studying BEST4^+^ epithelial cells, which are absent in mice. Solid food ingestion induced extensive transcriptome changes in each epithelial cell type, with the most pronounced changes noted in absorptive and BEST4^+^ cells. Some of the effects of solid food introduction were common to most epithelial cell types, such as the upregulation of *ALDH1A1*, which encodes for a vitamin A processing enzyme. Solid food ingestion remodeled epithelial defenses systems, as observed by the increased expression of interferon-stimulated genes in mature absorptive and BEST4^+^ cells. Solid food also upregulated the gene expression of the immunoglobulin transporter *PIGR*, specifically in cells located at the base of epithelial crypts and in goblet cells. In addition, solid food triggered epithelial differentiation, which was associated with modification of the expression of genes involved in handling of amino acids, lipids and bile acids, as well as changes in hormone expression by enteroendocrine cells. These cell type-specific transcriptome modifications induced by solid food ingestion coincided with changes in microbiota composition and metabolic activity, which may contribute to epithelial maturation. Overall, our work provides a single-cell atlas of the transcriptome changes induced in the intestinal epithelium at the suckling-to-weaning transition.

## Introduction

The intestinal epithelium contributes to digestion and allows nutrient absorption while providing a physical and immunological barrier against microorganisms and toxic compounds (Peterson and Artis 2014). This dual functionality is enabled by specialized absorptive and secretory epithelial cells, all derived from actively dividing stem cells located at the crypt base (Barker 2014). Single-cell transcriptomics has recently deepened our understanding of the cellular diversity of the intestinal epithelium (Elmentaite et al. 2020; Fawkner-Corbett et al. 2021; Burclaff et al. 2022). Absorptive cells now emerge as a heterogeneous population with distinct functions along the crypt-villus axis (Moor et al. 2018; Beumer et al. 2022). Additionally, BEST4^+^ cells were recently identified as a novel subset of mature absorptive cells with potential roles in ion transport, mucus hydration, and secretion of antimicrobial peptides and hormones (Malonga et al. 2024). Broad cellular diversity is also osberved in the secretory cell lineage, as exemplified by the numerous subtypes of enteroendocrine cells defined by their hormone secretion profiles (Beumer et al. 2020). Heterogeneous populations of mucus-secreting goblet cells have also been identified along the crypt-villus axis with a zonation of their antimicrobial activities (Parikh et al. 2019; Beumer et al. 2022). However, this newly uncovered diversity of epithelial cells is minimally understood in the context of postnatal intestinal development.

After birth, the intestinal epithelium of the newborn undergoes a maturation process that culminates at the suckling-to-weaning transition (Muncan et al. 2011; Frazer and Good 2022). Indeed, the dietary switch from maternal milk to solid food is associated with an adaptation of epithelial digestion, absorption and transport systems enabling the transition from a high-fat diet to a carbohydrate-based diet (Henning 1981; Hansson et al. 2011). The onset of solid food ingestion also coincides with a reduced epithelial permeability linked to a remodeling of epithelial defense systems including tight junctions, microbial detection systems, glycosylation and secretion of antimicrobial peptides and mucus (Biol-N’garagba and Louisot 2003; Holmes et al. 2006; Ménard et al. 2008; Price et al. 2018; Navis et al. 2019). This epithelial developmental process follows a genetically wired program tuned by several factors including changes in glucocorticoid levels, the introduction of solid food, and the cessation of suckling (Nanthakumar and Henning 1993; Muncan et al. 2011; Beaumont et al. 2022; Díez-Sánchez et al. 2024). In addition, ingestion of solid food also strongly alters the composition of the gut microbiota, which contributes to the induction of epithelial maturation, notably through the release of bacterial metabolites such as butyrate (Pan et al. 2018; Beaumont et al. 2020). A defect in host-microbiota co-maturation around weaning is known to increase the susceptibility to inflammatory or metabolic diseases later in life (Al Nabhani et al. 2019; Hornef and Torow 2020). It is therefore essential to understand the mechanisms underlying epithelial maturation at the suckling-to-weaning transition. However, a large knowledge gap exists regarding epithelial cell type-specific adaptations triggered by the introduction of solid food in the diet.

Transcriptomic regulations induced in the intestinal epithelium during the transition from milk to solid food were described in mice (Rakoff-Nahoum et al. 2015; Pan et al. 2018). However, these studies did not delineate the relative contributions of age and nutrition in epithelial maturation. In fact, controlling dietary intake in early life is difficult in mice because the stress of separation from the mother disrupts the gut barrier function (Ilchmann-Diounou and Menard 2020). In contrast, in rabbits, suckling occurs only once per day for about 5 minutes and there is naturally little contact between the mother and her litter (Gidenne and Lebas 2006). This unique behavior allows to control of milk and solid food ingestion in early life by housing separately the mother and her litter (Beaumont et al. 2022). In addition, we have recently shown that BEST4^+^ cells are present in the intestinal epithelium of rabbits, whereas these cells are absent in mice (Malonga et al. 2024). Thus, the rabbit is a valuable model to study epithelial maturation at the suckling-to-weaning transition. In this study, we used single-cell transcriptomics to identify the maturation program induced by solid food ingestion in each cell type of the intestinal epithelium of age-matched suckling rabbit littermates fed or not with solid food. In addition, analysis of the microbiota and metabolome allowed us to link the changes in luminal environment induced by solid food ingestion with the gene expression regulation observed at the single-cell level in the epithelium.

## Material and methods

### Animal experiments

The experiments were performed at the PECTOUL experimental facility (GenPhySE, INRAE, Toulouse, France). The handling of rabbits followed the recommendations outlined by the European Union’s regulations for the protection of animals used in scientific research (2010/63/EU), and was consistent with the French legislation (NOR: AGRG1238753A 2013). This project received approval from the local ethics committee “Comité d’éthique en expérimentation animale SCIENCE ET SANTE ANIMALES” N°115 (SSA_2022_012 and SSA_2024_004V2). Multiparous dams (n = 4) were housed individually in wire cages (61 × 68 × 50 cm) equipped with a closed nest (39 × 27 × 50 cm). The litter size was limited to 10 pups per litter (Figure S1A). From postnatal day (PND) 4, pups were placed in a new cage, adjacent to their mother’s cage. At PND12, litter sizes were reduced to 6 pups in order to maximize milk ingestion. Pups from each litter were separated in 2 cages on either side of their mother’s cage (3 pups/cage) to form two groups. In the first group (Milk), the pups were exclusively suckling. In the second group (Milk+Solid), the pups were suckling while having *ad libitum* access to commercial solid food pellets (StabiGreen, Terrya). During the whole experiment, the dam and the pups were placed once a day for 5-10 minutes in the nest of the dam’s cage for suckling before returning to their respective cages. Coprophagia was prevented by removing feces dropped by the mother in the nest after each suckling. Individual milk intake was quantified daily (from PND12 onwards) by weighing pups before and after suckling. Solid feed intake was measured daily at the cage level (3 pups) by weighing the feeder. The experiment was repeated a second time independently with n = 5 litters in order to collect samples for RNA *in situ* hybridization, immunohistochemistry, electron microscopy and calprotectin measurements. All other measurements were performed on samples collected during the first experiment.

### Sample collection

One male pup per litter (n = 4) and per group (Milk or Milk+Solid) was sacrificed after suckling at PND24 or PND25 by electronarcosis followed by exsanguination. Blood collected in EDTA tubes was centrifuged (1000 × g, 10 min, 4°C) and plasma was stored at −20°C. The caecum with its content and appendix was isolated and weighed. The content of the caecum was collected and kept at −80°C until microbiota and metabolome analysis. A fragment of caecal tissue was collected and placed in cold PBS without Ca^2+^/Mg^2+^ (ThermoFisher scientific, cat#10010-015) for epithelial cell isolation. Other sections of caecal tissue were fixed in i) Carnoy solution (60% ethanol, 30% chloroform, 10% glacial acetic acid) for 3 hours before transfer in 70% ethanol (samples used for Alcian Blue and Periodic Acid of Schiff staining), or in ii) 10% neutral-buffered formalin for 24 hours before transfer in 70% ethanol (samples used for immunohistochemistry and RNA *in situ* hybridization), or in iii) 0.1 M Sörensen phosphate buffer (pH 7.4) with 2% glutaraldehyde at 4°C (samples used for electron microscopy).

### Caecal epithelial cell isolation

Caecal tissue was opened longitudinally and washed with cold PBS to remove all content. The tissue was minced into 1 cm^2^ sections and washed with cold PBS. Tissue segments were transferred to 5 mL of a pre-warmed (37°C) digestion solution prepared in HBSS without Ca^2+^/Mg^2+^ (ThermoFisher Scientific, cat#14175095) and supplemented with 5 mM EDTA (ThermoFisher Scientific, cat#AM9260G) and 1 mM DTT (Sigma, cat# 10197777001). After incubation (20 minutes at 37°C under slow agitation at 15 rpm), epithelial crypts were detached by vigorous manual shaking for one minute. The crypt solution was then filtered (100 µm) before centrifugation (300 × g for 5 minutes at 4°C). The crypt pellet was resuspended in 10 mL of pre-warmed dissociation solution containing TrypLE (ThermoFisher, cat# 1205036) supplemented with 1 mg/mL DNAse I (Sigma, cat # 10104159001), 5 mM MgCl_2_ (Sigma, cat# M1028), 10 µM Y27632 (StemCell Technologies, cat# 72304) and the solution was distributed in two 50 mL conical tubes (5 mL/tube). Cells were incubated for 10 minutes at 37°C under gentle agitation at 15 rpm before homogenization by vortexing (3 seconds). This step was repeated and followed by two successive filtrations (70 µm and 40 µm). Digestion was stopped by adding 45 mL of cold PBS to the cells. After centrifugation (300 × g for 5 minutes at 4°C), the cells were resuspended in 5 mL FACS buffer (PBS supplemented with 3% fetal bovine serum [ThermoFisher Scientific, cat#10270-106], 2 mM EDTA, and 10 µM Y27632). Cell concentration was measured using an automated cell counter Countess 3 (ThermoFisher Scientific, cat#AMQAX2000).

### Cell preparation for single-cell sequencing

Cells (2×10^6^) were centrifuged (300 × *g* for 5 minutes at 4°C) and resuspended in 1 mL of PBS supplemented with 10 µM Y27632. This step was repeated twice. Dead cells were stained with the LIVE/DEAD™ Fixable Violet Dead Cell Stain Kit (ThermoFisher Scientific, cat#L34963), according to the manufacturer’s instructions. After 30 minutes of incubation (4°C, protected from light), cells were centrifuged (300 × *g* for 5 minutes at 4°C) and resuspended in 1 mL FACS buffer. This step was repeated once. Cells were filtered (40 µM) and sorted (10^5^ live and single-cells) in a 1.5 mL tube containing 10 µL of PBS supplemented with 10 µM Y27632 by using a BD Influx cell sorter instrument with a 100 µm nozzle, under 20 psi at the I2MC Cytometry and Cell sorting TRI platform (Toulouse, France). After centrifugation (300 × *g* for 5 minutes at 4°C), cells were resuspended in 100 µL PBS, counted manually and their viability was verified by trypan blue staining.

### Single-cell sequencing

For single-cell RNA-sequencing, approximately 10,000 cells per sample were used for encapsulation into droplets using Chromium Next GEM Single-cell 3′ Reagent Kits v3.1 according to manufacturer’s protocol (10x Genomics CG000315 Rev E user guide). Briefly, after generation of Gel bead-in-EMulsions (GEMs) using Next GEM Chip G, GEMs were reverse transcribed in a C1000 Touch Thermal Cycler (BioRad) programmed at 53°C for 45 min, 85°C for 5 min, and held at 4°C. After reverse transcription, single-cell droplets were broken and cDNA was isolated and cleaned with Cleanup Mix containing DynaBeads (ThermoFisher Scientific). cDNA was then amplified with a C1000 Touch Thermal Cycler programmed at 98°C for 3 min, 12 cycles of (98°C for 15 s, 63°C for 20 s, 72°C for 1 min), 72°C for 1 min, and held at 4°C. Subsequently, approximately 250 ng of amplified cDNA was fragmented, end-repaired, A-tailed, index adaptor ligated, and cleaned with cleanup mix containing SPRIselect Reagent Kit (Beckman Coulter, cat# B23317) in between steps. Post-ligation product was amplified and indexed with a C1000 Touch Thermal Cycler programmed at 98°C for 45 s, 11 cycles of (98°C for 20 s, 54°C for 30 s, 72°C for 20 s), 72°C for 1 min, and held at 4°C. The sequencing-ready libraries were cleaned up with SPRIselect beads. 10x libraries were pooled and charged with 1% PhiX on one S1 lane of the NovaSeq 6000 instrument (Illumina) using the NovaSeq 6000 S1 Reagent Kit v1.5 (100 cycles), and the following sequencing parameters: 28 bp read 1 – 10 bp index 1 (i7) – 10 bp index 1 (i5) – 150 bp read 2. The S1 lane generated a total of 810×10^6^ raw reads.

### ScRNA-seq pre-processing, filtering, normalization and clustering

Cell Ranger Software (version 7.1.0, 10x Genomics) was used to align and quantify raw sequencing data using the rabbit reference genome (GCF_009806435.1_UM_NZW_1.0). A custom reference file was created using the Cell Ranger mkgtf command with “--attribute=gene_biotype:protein_coding and -- attribute=gene_biotype:lncRNA” parameters. The Cell Ranger mkref and count commands were used with default parameters.

Using R software (version 4.2.1), the Seurat (version 4.3.0) pipeline (Butler et al. 2018) was run for data preprocessing and analysis. Cells with less than 1,600 or more than 55,000 expressed genes were filtered out. Similarly, cells with a number of counts below 1,500, with a percentage of mitochondrial RNA above 25% or expressing more than 0.1% of counts from hematopoietic cell genes (*CD44, PTPRC, CD48*) were filtered out. The resulting data were normalized via the *NormalizeData* function of Seurat, with the *LogNormalize* method. The top 2,000 variable features were then extracted (based on a mean-variance trend as implemented in the *FindVariableFeatures* function of Seurat). After scaling data to unit variance, dimensionality reduction was carried out with Principal Components Analysis (50 PCs). Cell clustering was performed based on retained PCs and using the Leiden algorithm on a cell similarity graph, with a 0.5 resolution. Finally, clusters were visualized using the non-linear reduction dimensionality Uniform Manifold Approximation and Projection (UMAP) performed on PCA reduction (30 PCs).

### Cell type assignment

#### Marker genes

The marker list for each cluster was obtained using a Wilcoxon test as implemented in the Seurat function *FindMarkers*. A gene was declared a marker if its adjusted *p*-value < 0.05 (Bonferroni correction for multiple testing). The test results were further filtered to ensure a minimum log-Fold Change (logFC) of 0.25 between the tested cluster and the others. Only genes expressed in at least 25% of cells and over-expressed in the tested cluster (compared to the others) were considered for this analysis. Cell types were then manually assigned to clusters according to found markers, based on a comparison with known cell type markers (Parikh et al. 2019; Elmentaite et al. 2021; Burclaff et al. 2022; Wiarda et al. 2023).

#### Assessment of cluster validity with cell cycle score and crypt axis gene score

The cell cycle score was used to assign phases of the cell cycle to individual cells and assess the consistency between manually assigned cell types and expected cell cycle phase. The cell cycle score was computed with the Seurat function *CellCycleScoring*. In addition, the crypt axis gene (CAG) score of each cell was calculated via the *AddModulesScore* function by averaging the expression of genes previously defined as expressed in epithelial cells located at the crypt top (*PLAC8, CEACAM1, TSPAN1, DHRS9, KRT20, RHOC, PKIB, HPGD*) (Parikh et al. 2019).

#### Pseudo-time analysis

Considering that absorptive cell (AC) subsets are distinguishable from each other based on their differentiation states, we used a pseudo-time analysis to create AC sub-groups with the monocle3 package (version 3.1). The trajectory of cell types was obtained using the *learn_graph* function on the previously generated UMAP and the pseudo-time of each cluster was calculated based on their projection on the trajectory using the *order_cells* function. The trajectory root was set to be the stem cell cluster. Subsequently, three cell subgroups of AC were delineated based on the pseudo-time distribution, which was found to be trimodal: cells with a pseudo-time below 2.0 were classified as “Early AC”, those with a pseudo-time between 2.0 and 8.6 were classified as “Intermediate AC”, and cells with a pseudo-time above 8.6 were classified as “Mature AC” (Figures S3A and B). Marker extraction was performed for each assigned cell type, similarly to what was performed for cluster markers and as described in the “Marker genes” section.

#### Automatic assignation of cell types

To validate our manual annotation, we performed an automatic annotation based on the transfer of labels of a reference to the rabbit caecum epithelial cells (Stuart et al. 2019). Human epithelial cells from the large intestine (Elmentaite et al. 2021) were used as a reference. First, a PCA was performed on the reference dataset to reduce its dimension to the first 30 PCs. Then, *FindTransferAnchors* was used to find similar cells between the rabbit and human datasets, called “anchors”. These anchors were then used by the *MapQuery* function to map the rabbit caecum epithelial cells onto the human epithelial cell space. The reference annotation was then transferred from the reference to the rabbit data and visualized on the UMAP. The results of *MapQuery* were also used in the *MappingScore* function to attribute a score to each rabbit cell. Roughly, this score measures how a cell neighborhood is affected by a mapping to and then back from the reference (a higher score corresponds to a more similar neighborhood).

#### Biological pathways enrichment

Biological enrichment analysis was performed on marker genes of the cell types using the *enrichGO* function from the *clusterProfiler* (version 4.6.1) package (Yu et al. 2012), with all expressed genes as the reference background. The enrichment analysis was carried out using the *Homo sapiens* database because of the absence of an *Oryctolagus cuniculus* database. Redundancy of results was reduced by using the *simplify* function from the *clusterProfiler* package. Terms with a semantic similarity over 0.7 were deleted and only representative terms (terms with the smallest *p*-value) were kept within each group of term. *p-*values were corrected for multiple testing using the Benjamini-Hochberg (BH) procedure (Benjamini and Hochberg 1995) and pathways were considered enriched if their corresponding adjusted *p*-value was < 0.05.

#### Differential analysis of gene expression

The pseudo-counts data were derived by summing the counts of each gene across cells of the same type for each rabbit. This step is considered essential as it has been shown that performing the differential analysis on pseudo-bulk data yields more robust results, reducing the risk of Type I errors compared to analyzing scRNA-seq data directly (Zimmerman et al. 2021; Murphy and Skene 2022). The whole analysis was performed independently in each cell type. Pseudo-counts were normalized using the “TMM” method of e*dgeR* (Robinson et al. 2010). A PCA was conducted on log2-transformed pseudo-counts for quality control, revealing a possible important impact of the litter on gene expression (Figure S4). This was thus accounted for in the differential analysis. Differential expression analysis was performed using a Negative Binomial generalized linear model as implemented in *edgeR*. More precisely, each gene expression was modeled with an additive effect of both the group and the litter, the latter being used as a blocking variable. *p-*values were obtained with a log-likelihood ratio (LR) test of the group effect. Adjusted *p*-values were obtained with the BH procedure and genes were considered differentially expressed if their corresponding adjusted *p*-value was < 0.05. Differentially expressed genes were subjected to an enrichment analysis as described in the *“*Biological pathways enrichment” section.

### Microbiota composition

The microbiota composition was analyzed as described previously (Beaumont et al. 2020). Briefly, DNA was extracted from 50 mg of caecal content with the Quick-DNA Fecal/Soil Microbe DNA Miniprep Kit (Zymo Research, cat#D6010). The V3-V4 region of the 16S gene was amplified by PCR and amplicons were sequenced by MiSeq Illumina Sequencing. Bioinformatic analyses were performed with the FROGS pipeline (v.4.0.1) according to the guidelines (Escudié et al. 2018). Taxa representing more than 0.5% of the relative abundance in at least one group were considered for analysis, as it was previously shown that taxa below this threshold were not accurately quantified (Paës et al. 2020).

### Metabolomics

The metabolome was analyzed in 50 mg of caecal content by using nuclear magnetic resonance (NMR)-based metabolomics, at the MetaboHUBMetaToul-AXIOM metabolomics platform (GenotToul, Toulouse, France), as described previously (Beaumont et al. 2020). The relative concentration of each metabolite was expressed relatively to the mean quantification measured in the Milk group.

### Plasma biochemistry

The Clinical Chemistry Analyzer Pentra C400 (Horiba medical) was used at the Anexplo Phenotyping platform (GenoToul, Toulouse) to measure plasmatic concentrations of cholesterol, high density lipoprotein (HDL), low density lipoprotein (LDL), glucose, triglycerides, free fatty acids and urea.

### Calprotectin assay

Caecal epithelial cells isolated as described above were lysed in RIPA buffer (ThermoFisher Scientific, cat#89901) supplemented with cOmplete protease inhibitor cocktail (Roche, cat#11697498001) by using stainless steel beads and a TissueLyser II (Qiagen) operating at 30 Hz for 3 min. Lysates were centrifuged (12000 × g, 10 min, 4°C) and stored at −80°C until analysis. Calprotectin was quantified in undiluted epithelial cell lysates by using a rabbit-specific ELISA kit (Clinisciences, cat# MBS2601529-48), following the manufacturer instructions. Protein concentration in epithelial cell lysates diluted 1:2 (v/v) in NaCl 0.9% was measured with Pierce Bradford Plus Protein Assay Kits (ThermoFisher Scientific, cat#23236). Calprotectin concentration was normalized to the protein level of each sample.

### Histology

Transversal sections of cecal tissue with luminal content fixed in Carnoy’s solution were embedded in paraffin and stained by Alcian Blue and Periodic Acid of Schiff at the histology platform Anexplo (GenoToul, Toulouse, France). Slides were digitalized before measurement of the crypt depth and of the number of goblet cells per crypt with the CaseViewer 2.3 software (3DHISTECH). Formalin-fixed, paraffin-embeded (FFPE) transversal sections of cecal tissues were cut into 4 µm sections and adhered to Superfrost-Plus charged microscope slides (Thermo Fisher Scientific) before being used for RNA *in situ* hybridization and immunohistochemical staining.

#### RNA in situ hybridization

The RNAscope 2.5 HD Reagent Kit – RED (Advanced Cell Diagnostics, cat#322350) and the RNAscope Multiplex Fluorescent Reagent Kit v2 (Advanced Cell Diagnostics, cat#323100) were used with rabbit custom probes targeting *SPINK4* (Advanced Cell Diagnostics, cat#1564251-C1, RNAscope™ Probe-Oc-SPINK4-C1) or *BEST4* (Advanced Cell Diagnostics, cat#1564261-C1, RNAscope™ Probe-Oc-BEST4-C1) or *PIGR* (Advanced Cell Diagnostics, cat#1003001-C1, RNAscope™ Probe-Oc-PIGR-C1). Negative and positive control slides were respectively hybridized with the RNAscope™ Negative Control Probe-DapB (Advanced Cell Diagnostics, cat#310043) and RNAscope™ Probe-Oc-GAPDH-No-XHs (Advanced Cell Diagnostics, cat# 469461) or RNAscope™ Probe-Oc-POLR2A (Advanced Cell Diagnostics, cat# 410571).

The chromogenic assay (*SPINK4*) was performed as described before with minor modifications (Palmer et al. 2019). Briefly, slides were incubated at 60°C to enhance tissue adherence. Slides were deparaffinized using xylene and rehydrated through a series of graded ethanol washes. Slides were treated with hydrogen peroxide to block endogenous peroxidase activity, followed by epitope unmasking using a boiling target retrieval solution. Hydrophobic barriers were drawn around the tissues to contain reagents. Unless stated otherwise, all incubations were performed at 40°C in a HybEZ hybridization oven followed by a 1x Wash Buffer wash. Slides were incubated with the Protease Plus for 15 minutes to break down RNA-associated proteins. Probes were then applied to each slide and incubated for 2 hours. The amplification steps were carried out sequentially using AMP1, AMP2, AMP3, AMP4, AMP5, and AMP6, with varying incubation times (30, 15, 30, 15, 30, and 15 minutes) and temperatures (40°C for AMP1–4, and room temperature for AMP5 and AMP6). Chromogenic detection was performed using a 1:60 dilution of RED-A:RED-B, followed by a counterstaining with Gill’s Hematoxylin I (American Master Tech Scientific, cat# HXGHE1LT, diluted 1:1 in dH O). Slides were immediately air-dried for 20 minutes at room temperature, after which two drops of Vecta Mount (Vector Laboratories, cat# H-5700-60) were applied. Coverslips (#1 thickness) (Fisherbrand, cat# 12-545-F) were mounted, and the slides were left to dry for 20 minutes at room temperature.

The fluorescent *in situ* hybridization (*PIGR* and *BEST4*) assay was performed identically to the chromogenic assay for the pretreatment and the hybridization steps. The amplification was performed with AMP1 (30 minutes), AMP2 (15 minutes), and AMP3 (15 minutes). Detection was performed with horseradish peroxidase channel 1 (HRP-C1) for 15 minutes and washed. The Opal 570 Reagent fluorophore (Akoya Biosciences, cat#FP1488001KT, dilution 1:750 in RNAscope Multiplex TSA Buffer [Advanced Cell Diagnostics, cat# 322809]) was incubated on the slides for 30 minutes. After the wash, the horseradish peroxidase (HRP) blocker was added to the slides and incubated for 15 minutes. Similarly to the chromogenic assay, incubations were performed at 40°C in a HybEZ oven, followed by a wash performed twice with 1X Wash Buffer for 2 minutes at RT. All the slides were incubated with DAPI (Advanced Cell Diagnostics, cat# 323108) for 30 seconds at RT and mounted with 2 drops of ProLong Gold Antifade reagent (Invitrogen, cat# P36930) and covered with #1.5 thickness cover glass (Fisherbrand, cat# 12-545-F).

#### Chromogenic immunohistochemistry (IHC)

The IHC assay was performed as described before (Wiarda et al. 2020). Briefly, slides were incubated for 20 minutes at 60°C, deparaffinized in xylene, and rehydrated with ethanol and distilled water using a histological automaton (Leica Biosystem, cat#ST5020). Antigen retrieval was performed by submerging the slides in preheated distilled water, followed by incubation in 1X sodium citrate solution for 15 minutes at 95°C. Hydrophobic barriers were drawn around the sections, and slides were incubated with Dual Endogenous Enzyme Block (Dako, cat#S2003) for 10 minutes at RT, Protein Block (Dako, cat#X0909) for 20 minutes at RT, and primary antibody ([Goat Anti-Rabbit IgA, Abcam Limited, cat#ab97186, 1:3000], [Mouse anti-KI67, BD Biosciences, cat#AB_393778; 1:40], dilution in 1% bovine serum albumin [BSA] PBS, 4°C overnight) sequentially. Slides were then incubated with a secondary antibody ([ImmPRESS™ HRP Anti-Goat Ig, Vector, cat# MP-7405] or [HRP Labelled Polymer Anti-Mouse, Dako, cat#K400111-2]) for 30 minutes at RT, followed by DAB chromogen for 7 minutes at RT in the dark, and counterstained with 25% diluted Gill’s hematoxylin (American Master Tech Scientific, cat#HXGHE1LT) for 1 minute at RT. Slide washes were performed after each incubation using a 0.05% PBS-Tween solution. Slides were dehydrated in ethanol and Propar clearant (Anatech, cat#510) sequentially, mounted with Refrax Mounting Medium (Anatech, cat#711), and covered with #1 thickness cover glass (Fisherbrand, cat#12-545-F) using a histological automaton (Leica Biosystem, cat#ST5020). Negative controls were treated with 1% BSA in PBS without the primary or secondary antibodies.

### Electron microscopy

Electron microscopy analyses were performed in CMEAB (Toulouse, France). *Transmission electron microscopy (TEM):* Following fixation, samples were washed overnight in 0.2 M Sörensen phosphate buffer (pH 7.4). Post-fixation was carried out at room temperature for 1 hour in 0.05 M Sörensen phosphate buffer (pH 7.4) with 1% OsO_4_ and 0.25 M glucose. Dehydration was performed using graded ethanol series at room temperature, up to 70%. From then, the tissues were embedded in Embed 812 resin (Electron Microscopy Sciences) using a Leica EM AMW automated microwave tissue processor for electron microscopy. Once poylymerized, the samples were sliced into ultrathin sections (70 nm) using an Ultracut Reichert Jung ultramicrotome and mounted onto 100-mesh Formvar-coated copper grids. Sections were then stained with 3% uranyl acetate in 50% ethanol and Reynold’s lead citrate. Examinations were conducted on a transmission electron microscope (Hitachi HT7700) at an accelerating voltage of 80 kV. *Scanning electron microscopy (SEM):* After washing the sample in water, dehydration was performed through a graded ethanol series, up to 100% ethanol. Critical point drying was carried out with a Leica EM CPD 300. Dried samples were then coated with a 6 nm layer of platinum using a Leica EM MED020. SEM imaging was performed using a FEG FEI Quanta 250 scanning electron microscope at an accelerating voltage of 5 kV.

### Statistical analyses of microbiota, metabolites and calprotectin data

Statistical analyses of the log transformed relative abundances of bacterial taxa and metabolite concentrations (caecum or plasma), and calprotectin concentration in epithelial cells were analyzed with the R software (version 4.2.1). Linear mixed models were used with the group as a fixed effect and the litter as a random effect. *p-*values were corrected for multiple testing using the BH procedure. The significance of the group effect was tested and results were considered significant if their corresponding adjusted *p*-value was < 0.05.

## Data availability

The scRNA-seq data for this study have been deposited in the European Nucleotide Archive (ENA) at EMBL-EBI under accession number PRJEB74645 (www.ebi.ac.uk/ena/browser/view/PRJEB74645). The data are also accessible on the FAANG portal (www.data.faang.org/dataset/PRJEB74645) and are publicly available on the Broad Institute Single-cell Portal (https://singlecell.broadinstitute.org/single_cell/study/SCP2662/single-cell-transcriptomics-in-caecum-epithelial-cells-of-suckling-rabbits-with-or-without-access-to-solid-food). 16S sequencing data have been deposited in NCBI Sequence Read Archive (SRA) under accession number PRJNA1130383. NMR raw spectra have been deposited in Metabolights under accession number MTBLS10648.

## Results

To evaluate the effects of solid food introduction on epithelial maturation, we determined single-cell transcriptomic profiles of caecal epithelial cells isolated from age-matched littermate suckling rabbits ingesting or not solid food (Milk group: n = 4, Milk+Solid group: n = 4, Figure S1A). Growth and milk intake were similar in the two groups (Figures S1B and C). In the Milk+Solid group, the small amount of solid food ingested (< 25g/day/rabbit) increased dramatically the weight of the caecum (Figures S1D and E).

### A single-cell atlas of the rabbit gut epithelium

After applying quality filters and removal of hematopoietic cells, the dataset included 13,805 caecal epithelial cells (Figures S2A-E). We identified 13 clusters of cells based on their transcriptome profiles (Figures S2F-G). Each cluster was assigned to a cell type based on expression of known markers of epithelial lineages (Figures 1A-C, Figure 2A and Table S1), pseudo-time analysis (Figure 1D, Figure S3A and B) and cell cycle score (Figure 1E). The proportion of each cell type was calculated to determine if it was consistent with the literature (Figure 2C). Crypt position score (Figure 2B), enriched biological processes (Figure 2D, Table S2), and automatic annotation (Figure S3C) were used as quality controls for our manual annotation. Tables S1 and S2 respectively provide the lists of marker genes and of enriched biological processes, for each cell type.

**Figure 1.**
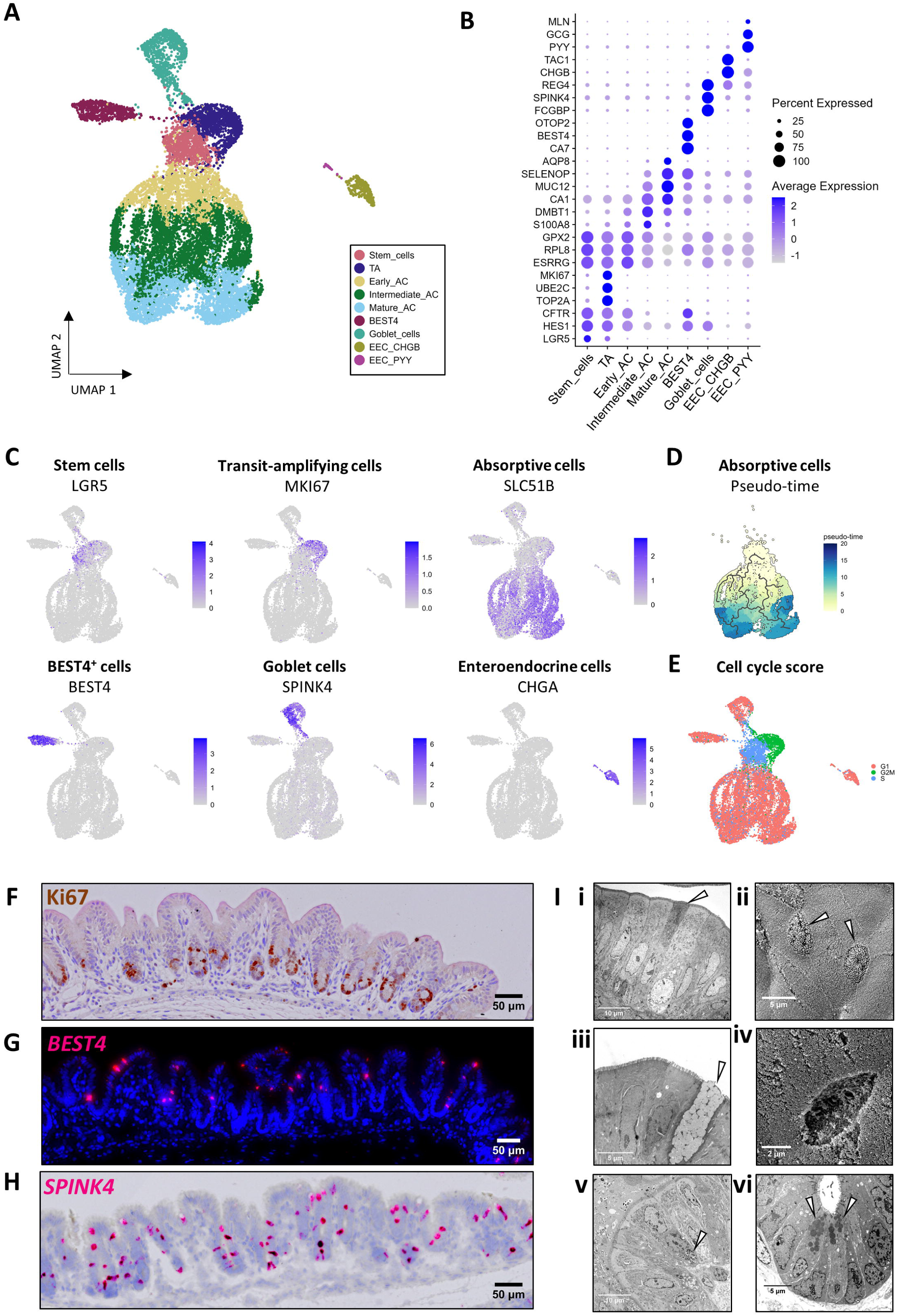
A single-cell transcriptomic atlas of the rabbit caecum epithelium. (A) Uniform Manifold Approximation and Projection (UMAP) of cells colored by epithelial cell type. The 13,805 cells were derived from n=8 suckling rabbits ingesting or not solid food (n=4/group). (B) Expression of selected marker genes for each cell type (average expression across cells in color and percentage of cells expressing the marker in size). (C) UMAPs colored by the expression of marker genes of each cell type. (D) UMAP colored by the pseudotime of the stem and absorptive cells. (E) UMAP colored by the inferred cell cycle state. (F) Localization of transit amplifying cells in the rabbit caecum epithelium by immunostaining of KI67 (brown). Nuclei are stained in blue. Scale bar 50 µm. (G) Localization of BEST4^+^ cells in the rabbit caecum epithelium *by in situ* hybridization of BEST4 mRNA (red). Nuclei are stained in blue. Scale bar 50 µm. (H) Localization of goblet cells in the rabbit caecum epithelium *by in situ* hybridization of SPINK4 mRNA (red). Nuclei are stained in blue. Scale bar 50 µm. (I) Ultrastructural appearance of rabbit caecum epithelial cells. i) Transmission electron microscopy (TEM) observation of absorptive cells. The white arrowhead shows an electron dense absorptive cell. Scale bar 10 µm. ii) Scanning electron microscopy (SEM) observation of microvilli. White arrowheads show cells with low density of microvilli. Scale bar 5 µm. iii) v) TEM observation of a goblet cell. The white arrowhead shows mucin granules. Scale bar 5 µm. iv) SEM observation of a goblet cell. Scale bar 2 µm. v) TEM observation of an enteroendocrine cell containing basal electron dense granules (white arrowhead). Scale bar 10 µm. vi) TEM observation of Paneth-like cells containing apical electron dense granules (white arrowheads) at the crypt base. Scale bar 5 µm AC: absorptive cells, EEC: enteroendocrine cells, TA: transit amplifying cells

**Figure 2.**
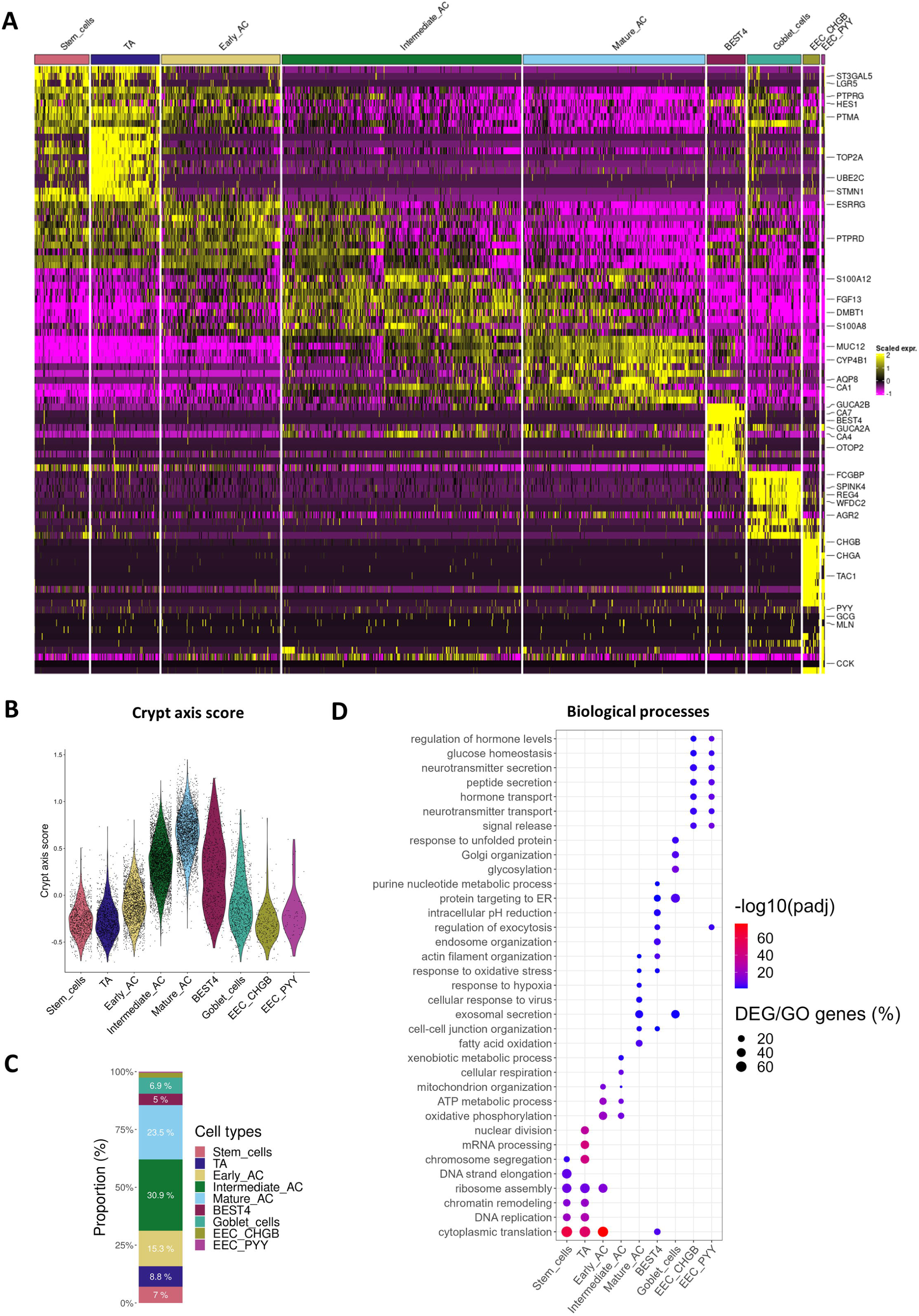
Transcriptionally distinct cell populations in the rabbit caecum epithelium. (A) Average expression of the top 10 marker genes with the highest average log2(fold change) for each cell type. For a given cell type, markers were ordered by decreasing log2(fold change) of the expression between this cell type and the other types. (B) Crypt axis score for each cell type. (C) Relative abundance for each cell type. (D) Selected biological processes enriched in marker genes for each cell type. The color corresponds to the -log10(adjusted p-value) value of the over-representation test and the size corresponds to the percentage of marker genes among the genes of the ontology term. AC: absorptive cells, EEC: enteroendocrine cells, TA: transit amplifying cells

Stem (*LGR5*^+^, 7% of epithelial cells) and transit amplifying (TA) cells (*MKI67^+^, TOP2A^+^, UBE2C*^+^, 8.8% of epithelial cells) were predicted to be positioned at the crypt base (Figures 1B-C, Figures 2A-C). Immunodetection of Ki67 (encoded by *MKI67*) confirmed that TA cells were localized at the crypt base (Figure 1F). Stem and TA cells were predicted in the S and G2M cell cycle phases, respectively, while most other epithelial cells were in the G1 phase (Figure 1E). Both stem and TA cells expressed high levels of genes involved in DNA replication and translation (Figure 2D). TA cells specifically expressed genes related to mRNA processing and nuclear division (Figure 2D). Absorptive cells (*SLC51B*^+^) were the main epithelial cell population and the distribution of their pseudo-times indicated three main differentiation states that we termed as early (15.3% of epithelial cells), intermediate (30.9% of epithelial cells) and mature (23.5% of epithelial cells) (Figures 1C-D, Figure 2C and Figures S3A-B). Absorptive cell gene expression profiles showed gradual modifications according to their differentiation states, which reflect their maturation during migration along the crypt axis (Figure 2A). Indeed, early absorptive cells were predicted to be localized at the lower part of crypts, while mature absorptive cells were positioned at the crypt-top (Figure 2B). Gene expression profile of early absorptive cells was intermediate between stem/TA cells and other absorptive cells (Figure 1B and Figure 2A). Intermediate absorptive cells expressed high levels of genes involved in antimicrobial defenses (e.g., *S100A8, S100A12, DMBT1*) and in mitochondrial metabolism (Figure 1B, Figures 2A and D, Table S1 and Table S2). Mature absorptive cells highly expressed genes involved in epithelial digestion, transport, and glycocalyx formation (e.g., *CA1, AQP8, ABCA1, APOB, ANPEP, MUC12*) (Figure 1B, Figure 2A and Table S1), and genes involved in lipid metabolism, response to hypoxia, and antimicrobial defenses (Figure 2D and Table S2).

BEST4^+^ cells (5% of epithelial cells), a recently discovered subset of mature absorptive epithelial cells, were identified by the expression of the canonical markers *BEST4*, *CA7, OTOP2, GUCA2B, GUCA2A,* and *CFTR* (Figures 1B and C, Figures 2A and 2C and Table S1). BEST4^+^ cells were predicted to be distributed along the crypt axis (Figure 2B). *BEST4* mRNA *in situ* hybridization confirmed that BEST4^+^ cells are relatively rare (< 4 BEST4^+^ cells per crypt) and distributed along the crypt axis (Figure 1G). Functions enriched in BEST4^+^ cells included “regulation of exocytosis” and “intracellular pH reduction” (Figure 2D and Table S2). Although the morphological features of BEST4^+^ cells are not defined yet, we observed rare electron dense absorptive cells and cells with low density microvilli that may correspond to distinct subsets of absorptive epithelial cells (Figures 1I-i and ii).

Goblet cells (6.9% of epithelial cells) were identified by expression of known markers of this lineage and of major components of mucus (*SPINK4, REG4, FCGBP, WFDC2, AGR2, ZG16, TFF3*) (Figures 1B and C, Figure 2A and C and Table S1). Goblet cells were predicted to be distributed across the crypt axis (Figure 2B) and we confirmed their localization by *SPINK4* mRNA *in situ* hybridization (Figure 1H). Genes specifically expressed by goblet cells were involved in “glycosylation” and “Golgi organization” (Figure 2D and Table S2), which reflects their role in the synthesis of mucins, also illustrated by goblet cells morphological features (Figures I-iii and iv). Enteroendocrine cells (EEC) (*CHGA^+^, NEUROD1*^+^) specifically expressed genes involved in the secretion of hormones (Figures 1C and Figure 2A, Tables S1 and S2). Two subclusters of EEC were distinguished based on their repertoire of hormone-related genes. EEC CHGB^+^ (2.2% of epithelial cells, enterochromaffin-like cells) expressed *TAC1*, *TTR, NMU*, and *TPH1* whereas EEC PYY^+^ (0.4% of epithelial cells, L-like cells) expressed *GCG*, *MLN*, and *CCK* (Figure 1B, Figure 2A and Table S1). Electron microscopy confirmed the presence of rare enteroendocrine cells containing electron dense granules at the basal side (Figure I-v).

Other rare cell types described in the intestinal epithelium of other species (tuft cells, Paneth cells, M cells) were not found in our rabbit gut scRNA-seq dataset. However, we observed at the base of epithelial crypts a few Paneth-like cells containing electron dense apical granules, whose scarcity and/or sensitivity to microfluidic flow may preclude their capture in droplets (Figure 1I-vi). Automatic cell annotation based on human large intestine scRNA-seq data was consistent with the manual assignment of cell types (Figure S3C). The mapping score indicating the degree of similarity between rabbit and human cells was the highest for stem cells, TA cells, BEST4^+^ cells, mature absorptive cells and for subsets of goblet and enteroendocrine cells (Figure S3D). All cell types were identified in each rabbit (Figure S3E). In sum, our analysis provided the first single-cell transcriptomic atlas of the rabbit intestinal epithelium. We have made these gene expression data available as a searchable tool on the Broad Institute Single-cell Portal (https://singlecell.broadinstitute.org/single_cell/study/SCP2662/single-cell-transcriptomics-in-caecum-epithelial-cells-of-suckling-rabbits-with-or-without-access-to-solid-food#study-visualize).

### Solid food introduction induced both global and cell type specific transcriptomic modifications in the intestinal epithelium

After characterizing the cellular diversity of the rabbit intestinal epithelium, we focused our analysis on the effects of solid food introduction on gene expression in each epithelial cell type. Ingestion of solid food altered the transcriptome of absorptive cells, as suggested in the UMAP by the low overlap between absorptive cells from suckling rabbits ingesting or not solid food (Figure 3A). Accordingly, the highest number of differentially expressed genes (DEG) was found in intermediate and mature absorptive cells with 890 and 868 DEG, respectively (Figures 3B-D, Figure S4). Although to a lesser extent, solid food introduction also modified the transcriptome in BEST4^+^ cells (429 DEG), early absorptive cells (268 DEG), TA cells (209 DEG), goblet cells (198 DEG), stem cells (189 DEG), EEC PYY^+^ (54 DEG), and EEC CHGB^+^ (41 DEG) (Figures 3B-D). These solid food induced alterations of gene expression were observed despite the proportion of epithelial cell types remaining similar in the two groups (Figure 3C). Table S3 provides the list of DEGs for each cell type. Table S4 contains the results of the enrichment analysis using DEGs of each cell type. All the biological functions and genes cited below were significantly modulated following the introduction of solid food (adjusted *p*-value < 0.05).

**Figure 3.**
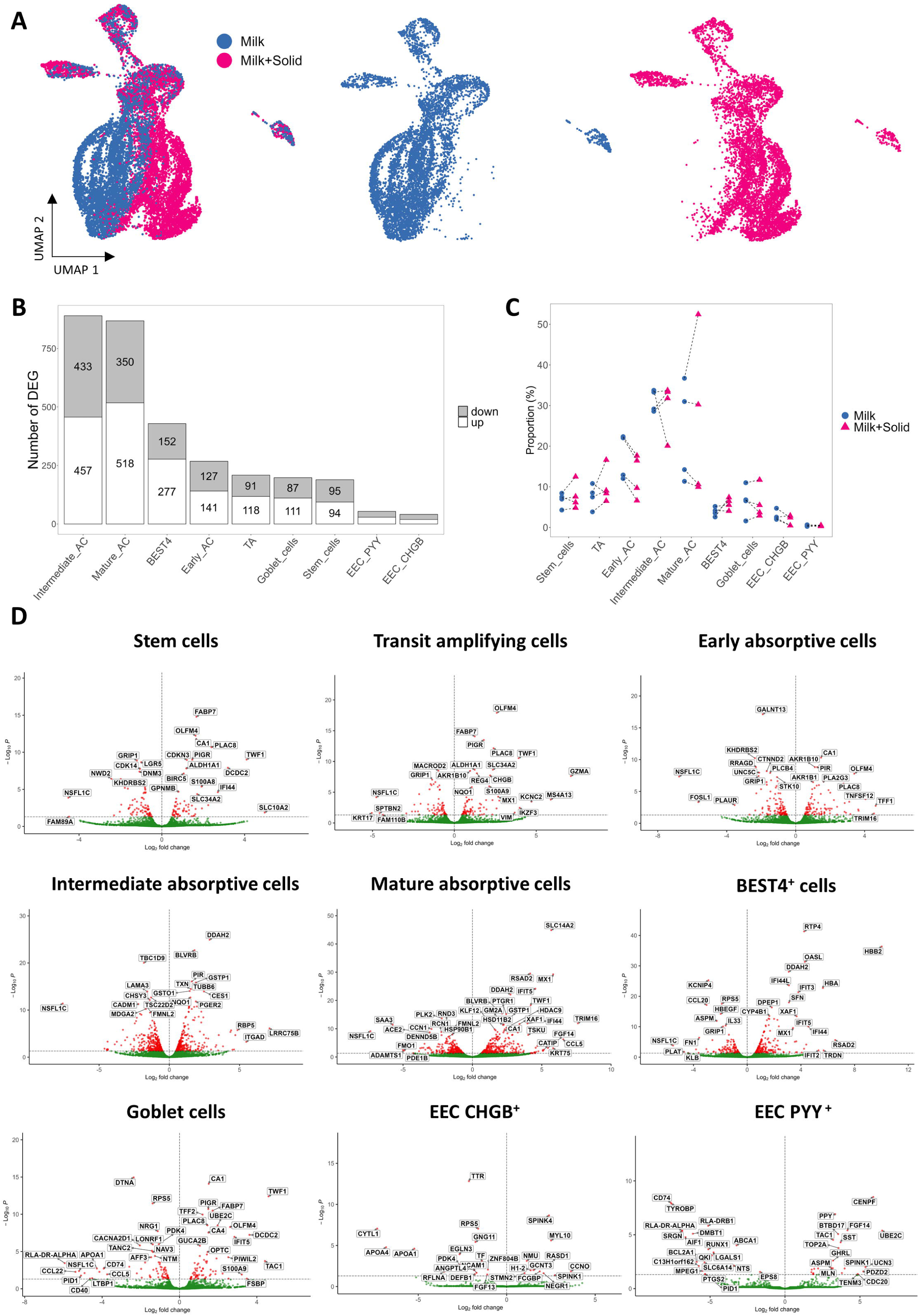
Ingestion of solid food by suckling rabbits modulates the transcriptome of each epithelial cell type. (A) Uniform Manifold Approximation and Projection (UMAP) of epithelial cells colored by group. The left panel shows UMAP of merged datasets, with cells restricted to Milk (n=4) and Milk+Solid (n=4) groups shown independently on the middle and right panels, respectively. (B) Number of differentially expressed genes (DEGs) between groups per cell type. Gray bars represent downregulated genes in the “Milk+Solid” group while white bars represent upregulated genes in the “Milk+Solid” group. DEG were obtained by using Negative Binomial generalized linear models on pseudo-bulk data fitted independently in each cell type. (C) Relative abundance of each epithelial cell type. Points represent individual values in rabbits and dotted lines link littermates. (D) Volcano plots of test results for each cell type. The -log10(adjusted p-value) values are plotted on the y-axis and the log2(fold change) values are plotted on the x-axis. AC: absorptive cells, EEC: enteroendocrine cells, TA: transit amplifying cells

Most of the modifications of gene expression induced by the introduction of solid food were cell type specific while other changes were shared between cell types (Figure 3D). Notably, mature absorptive cells and BEST4^+^ cells shared a high number of DEG (Figure 4A). Among transcriptomic modifications shared between most cell types, solid food introduction induced a strong upregulation of *ALDH1A1*, encoding an enzyme involved in retinoic acid metabolism, and an upregulation of *CA1*, a typical marker of epithelial differentiation in the large intestine (Figures 4B-D). Solid food introduction also increased the gene expression of the immunoglobulin transporter *PIGR* in several cell types, and this effect was much more pronounced in cells located at the bottom of the crypts (Figures 4B and E). *PIGR* mRNA *in situ* hybridization confirmed its predominant expression at the base of epithelial crypts (Figure 4F). The expression of *PIGR* was also increased by solid food ingestion in goblet cells that were found to contain immunoglobulin A (IgA) (Figures 4E and G). Among the cell type specific modifications, solid food ingestion reduced the gene expression of *LRIG1*, a master regulator of the stem cell niche, exclusively in stem cells (Figure 4B). Conversely, ingestion of solid food upregulated the gene expression of the transcellular water transporter *AQP8*, mostly in mature absorptive cells (Figure 4B and H). In BEST4^+^ cells, solid food introduction increased the expression of the pH-sensitive ion channel *OTOP2*, while it reduced the expression of the interleukin *IL33* (Figures 4B, I, and J). Overall, our results showed that the introduction of solid food induced major transcriptomic modifications in the intestinal epithelium of suckling rabbits and these changes are either shared across cell types or cell type specific. Accordingly, enrichment analyses revealed that solid food ingestion altered specific functions in every epithelial cell type (Figure 4K and Table S4).

**Figure 4.**
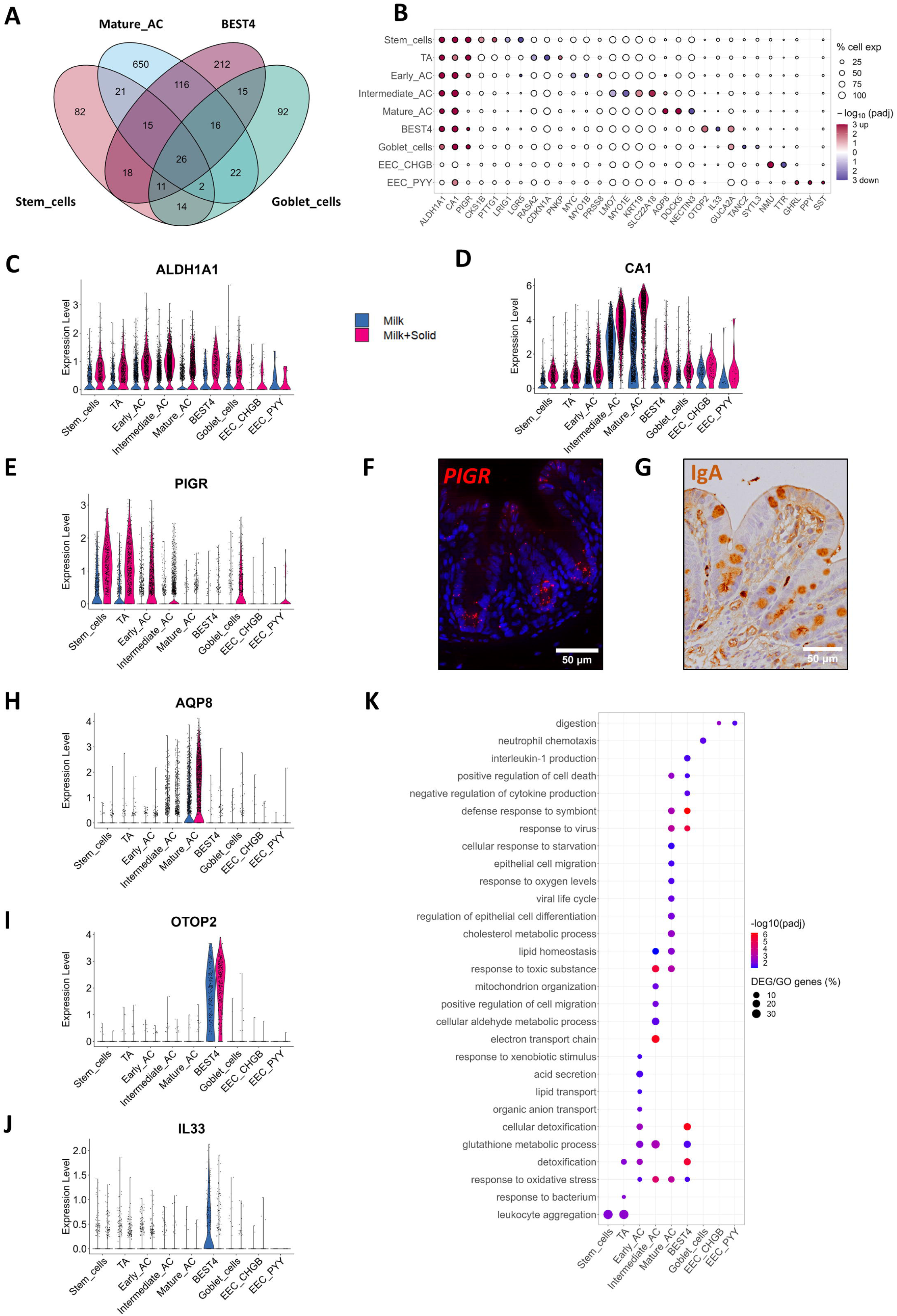
The transcriptomic changes induced by the introduction of solid food are predominantly cell-type specific. (A) Number of differentially expressed genes (DEGs) that are cell type-specific or shared by multiple cell types for stem cells, mature absorptive cells, BEST4^+^ cells, and goblet cells. (B) Selected DEGs significance (log10(adjusted p-value), color) and fold change sign (red for over expressed and blue for under expressed in the “Milk+Solid” group versus the “Milk” group) by cell type. The size represents the percentage of cells expressing the gene in the corresponding cell type. (C-E) Expression level of selected DEGs shared between several cell types, by cell (dot) per cell type and group. (F) Localization of PIGR mRNA (red) by *in situ* hybridization in the rabbit caecum epithelium. Nuclei are stained in blue. Scale bar 50 µm. (G) Immunostaining of immunoglobulin A (IgA, brown) in the rabbit caecum epithelium. Nuclei are stained in blue. Scale bar 50 µm. (H-J) Expression level of selected DEGs specific of one cell type, by cell (dot) per cell type and group. (K) Selected biological processes enriched in DEGs of each cell type. The color corresponds to the - log10(adjusted p-value) values and the size represents the percentage of DEGs included in the biological process. AC: absorptive cells, EEC: enteroendocrine cells, TA: transit amplifying cells

### Solid food introduction remodels defense systems in the intestinal epithelium

Solid food introduction upregulated the expression of genes involved in detoxification in all cell types, except EEC (e.g., *GPX2, GSTO1, GSTP1, MGST1, MGST3, SOD1, TXN*) (Figure 5A). This was particularly marked in intermediate and mature absorptive cells, and in BEST4^+^ cells. This finding is linked to the enrichment of biological pathways related to “cellular aldehyde metabolic process”, “response to toxic substance”, and “response to oxidative stress” in absorptive cells and in BEST4^+^ cells (Figure 4K). Solid food ingestion also increased the expression of interferon-stimulated genes (ISG), primarily in mature absorptive cells and BEST4^+^ cells (e.g., *DHX58, OASL, IFIT3, IFI35, IFI44L, IRF9, MX1, USP18, RIGI*) (Figure 5B), which is consistent with the specific enrichment of biological pathways such as “response to virus” and “defense response to symbiont” in these cell types (Figure 4K). Conversely, solid food ingestion decreased the expression of several genes coding for regulators of innate immune responses in absorptive cells (e.g., *AREG, NFKBIA, NFKBIZ*) (Figure 5C). This was associated with a cell type-specific downregulation of genes coding for cytokines in BEST4^+^ cells (*CXCL9, IL13RA1, IL33*), which were also characterized by a specific enrichment of the biological pathway “negative regulation of cytokine production” (Figure 4K). Other cell type specific downregulations of cytokine gene expression induced by solid food ingestion included *IL1A* in mature absorptive cells and *CCL25* in stem and early absorptive cells. In contrast, solid food introduction upregulated the expression of other cytokines expressed by small subsets of absorptive and BEST4^+^ cells (e.g., *IL18, IL32, IL34*) (Figure 5C). Solid food reduced the expression of some antimicrobial peptides in mature absorptive cells (*DMBT1* and *DEFB1*) and in goblet cells (*WDFC2*) while increasing the expression of numerous antimicrobial proteins of the S100 family in several cell types (e.g., *S100A1, S100A12, S100A14, S100A6, S100G*) (Figure 5D). Interestingly, genes coding for the two subunits of the inflammation marker calprotectin (*S100A8/S100A9*) were upregulated, notably in subsets of stem and TA cells (Figure 5D). The increased expression of calprotectin by epithelial cells after the ingestion of solid food was confirmed at the protein level in an independent experiment (Figure 5E).

**Figure 5.**
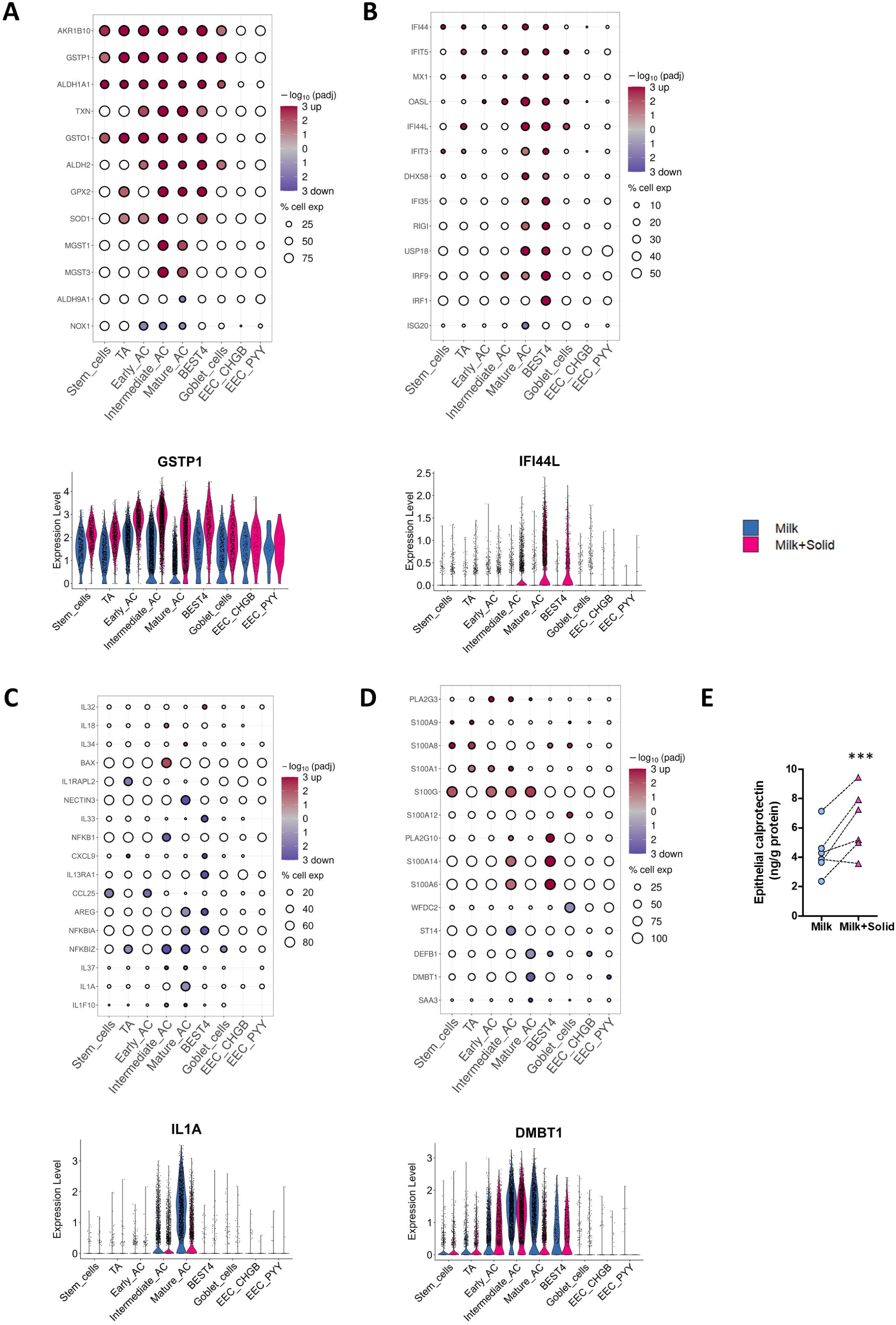
The introduction of solid food modifies the expression of genes involved in epithelial defenses. (A-D) Top: Selected differentially expressed genes (DEG) significance (log10(adjusted p-value) values, color) and fold change sign (red for over expressed and blue for under expressed genes in the “Milk+Solid” group versus the “Milk” group) involved in (A) detoxification and redox balance, (B) interferon signaling, (C) cytokine signaling, (D) antimicrobial peptides. The size corresponds to the percentage of cells expressing the gene in the cell type. Bottom: Expression level of a selected DEG by cell (dot), per cell type and group. (E) Concentration of calprotectin in caecum epithelial cells. Points represent individual values per rabbit and dotted lines link littermates. ***: p-value < 0.001. AC: absorptive cells, EEC: enteroendocrine cells, TA: transit amplifying cells

Solid food introduction also modulated the expression of numerous genes involved in epithelial glycosylation, which plays a key role in host-microbiota interaction. Solid food decreased the expression of several genes coding for glycosyltransferases mostly in stem, TA, early absorptive and goblet cells (e.g., *GALNT13, GALNT18 ST3GAL5, ST6GAL1, ST6GAL2*) while *B4GALT1* was upregulated in mature absorptive and BEST4^+^ cells (Figure 6A). In contrast, the introduction of solid food increased the expression of genes coding for fucosyltransferases (*FUT2* and *FUT9*) which are expressed by subsets of absorptive cells (Figure 6A). The introduction of solid food also altered the expression of genes related to mucin production (Figure 6B). Specifically, solid food ingestion reduced the expression of genes encoding the glycocalyx-forming transmembrane mucins *MUC1* and *MUC13* in intermediate and mature absorptive cells, while enhancing *MUC12* expression in BEST4^+^ cells (Figure 6B). In goblet cells, the expression of genes coding for major mucus components were upregulated (e.g., *TFF1, TFF2, ZG16*) or downregulated (e.g., *BCAS1, SYTL2*) after the introduction of solid food (Figure 6B). Histological observations confirmed that the number of goblet cells per crypt was similar in the caecal epithelium of rabbits ingesting or not solid food (Figures 6C-D), which is consistent with the goblet cell proportion estimation obtained by scRNA-seq (Figure 3C).

**Figure 6.**
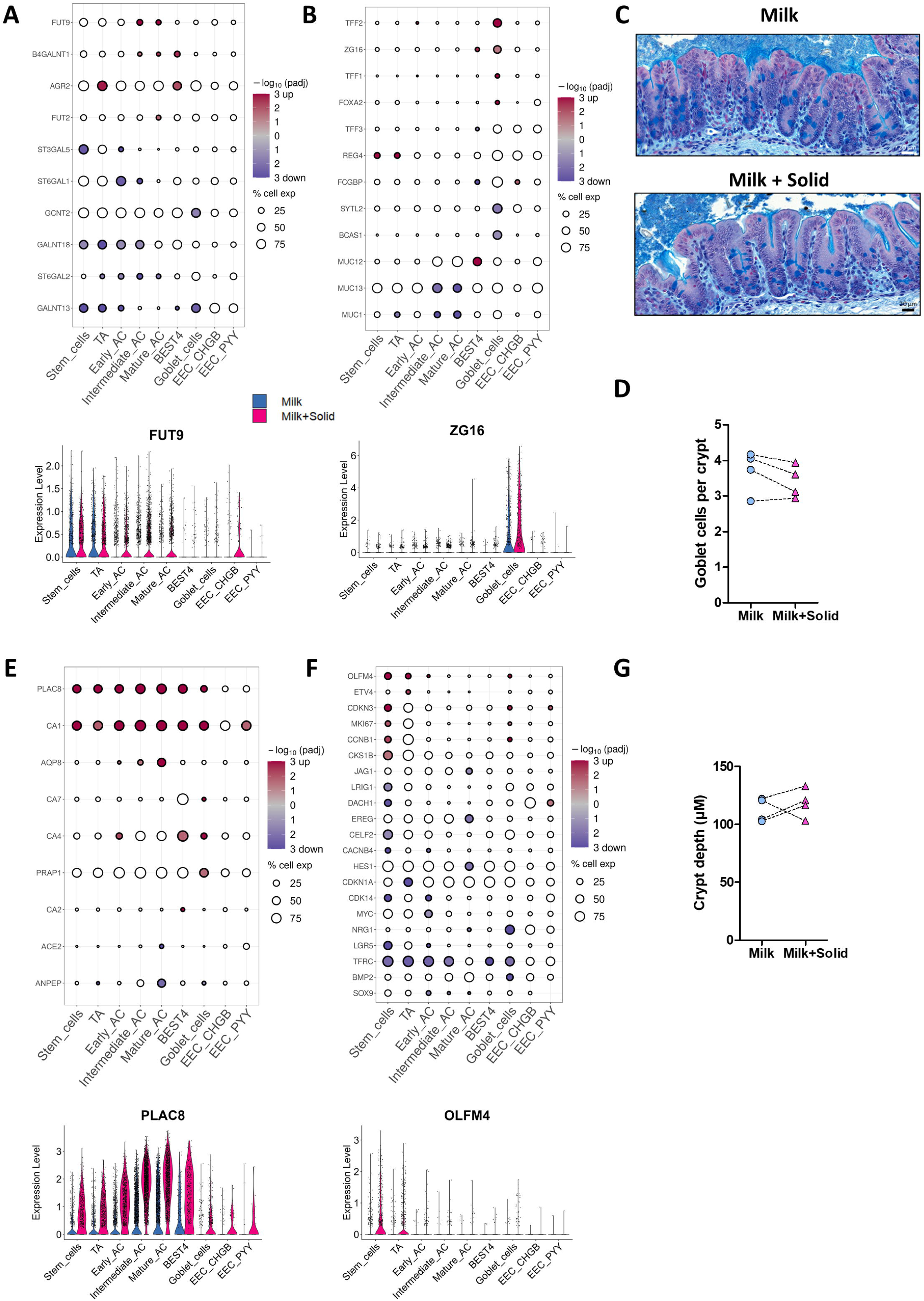
The introduction of solid food modifies the expression of genes involved in the mucus barrier, epithelial differentiation and renewal. (A-B) Top: Selected differentially expressed genes (DEG) significance (log10(adjusted p-value) values, color) and fold change sign (red for over expressed and blue for under expressed genes in the “Milk+Solid” group versus the “Milk” group) involved in (A) glycosylation, and (B) mucus components. The size corresponds to the percentage of cells expressing the gene in the cell type. Bottom: Expression level of a selected DEG by cell (dot), per cell type and group. (C) Representative histological observation of the caecum mucosa in each group. Alcian blue shows acidic mucins. Tissues are counterstained with hematoxylin and eosin. Scale bar 20 µm. (D) Number of goblet cells per crypt. Points represent individual values in rabbits and dotted lines link littermates. (E-F) Top: Selected DEG significance (log10(adjusted p-value) values, color) and fold change sign (red for over expressed and blue for under expressed genes in the “Milk+Solid” group versus the “Milk” group) involved in (E) differentiation, and (F) stemness and proliferation. The size corresponds to the percentage of cells expressing the gene in the cell type. Bottom: Expression level of a selected DEG by cell (dot), per cell type and group. (G) Epithelial crypt depth. Points represent individual values in rabbits and dotted lines link littermates. AC: absorptive cells, EEC: enteroendocrine cells, TA: transit amplifying cells

In order to determine whether changes in the expression of genes involved in epithelial defenses were linked to a change in the microbiota, we performed 16S rRNA gene sequencing in rabbit caecal contents (Table S5). Solid food ingestion by suckling rabbits altered the composition of the microbiota, in particular by increasing the abundance of the *Lachnospiraceae* family (14.6% in the Milk group versus 21.2% in the Milk+Solid group). Altogether, our results show that the introduction of solid food triggered major adaptations of epithelial defense systems in most cell types, which were associated with an alteration of the microbiota composition.

### Solid food introduction enhances differentiation in intestinal epithelial cells and alters nutrient handling

As a next step, we evaluated how solid food ingestion altered the expression of genes related to epithelial differentiation (Figure 6E) and renewal (Figure 6F). Key genes involved in stemness and proliferation were upregulated (e.g., *CDKN3, CKS1B, MKI67, OLFM4*) or downregulated (e.g.*, CDK14, CELF2, DACH1, LGR5, LRIG1*) in stem cells after the introduction of solid food (Figure 6F). These changes at the gene expression level were not associated with a modification of the crypt depth, which is partly determined by epithelial proliferation rate (Figures 6C and G). In contrast, solid food ingestion strongly increased the expression of the differentiation markers *PLAC8* and *CA1* in most of epithelial cells (Figure 6E). Moreover, the solid food induced upregulation of *AQP8* in mature absorptive cells was associated with the enrichment of the biological pathway “regulation of epithelial cell differentiation” (Figures 4K and 6E). Conversely, in mature absorptive cells, solid food introduction downregulated the expression of *ANPEP*, an enzyme involved in peptide digestion, which is a process usually occurring in the small intestine (Figure 6E). Accordingly, automatic annotation of cell types identified a population of enterocyte-like cells in the group of exclusively suckling rabbits (Figure S3C).

These results suggesting a rewiring of absorptive cell functions after solid food introduction were associated with the modulation of the expression of numerous genes involved in lipid handling and chylomicron biogenesis, mostly in absorptive cells (Figure 7A). These genes were either downregulated (*ABCA1, APOB, PLIN2, VLDR*) or upregulated (*ACAT2*, *APOM, LDLR)* (Figure 7A). Accordingly, absorptive cell DEGs were enriched in functions related to “lipid transport”, “lipid homeostasis”, and “cholesterol metabolic process” (Figure 4K). Moreover, solid food ingestion increased the expression of several bile acid transporters, such as *FABP7* that was upregulated in most cell types, *FABP6* that was specifically upregulated in BEST4^+^ cells, and *SLC51A* and *SLC51B* that were mostly upregulated in absorptive cells (Figures 7A and B). These alterations of the expression of lipid processing genes were coupled with an important decrease in the plasmatic concentration cholesterol and LDL after solid food introduction (Figure 7D).

**Figure 7.**
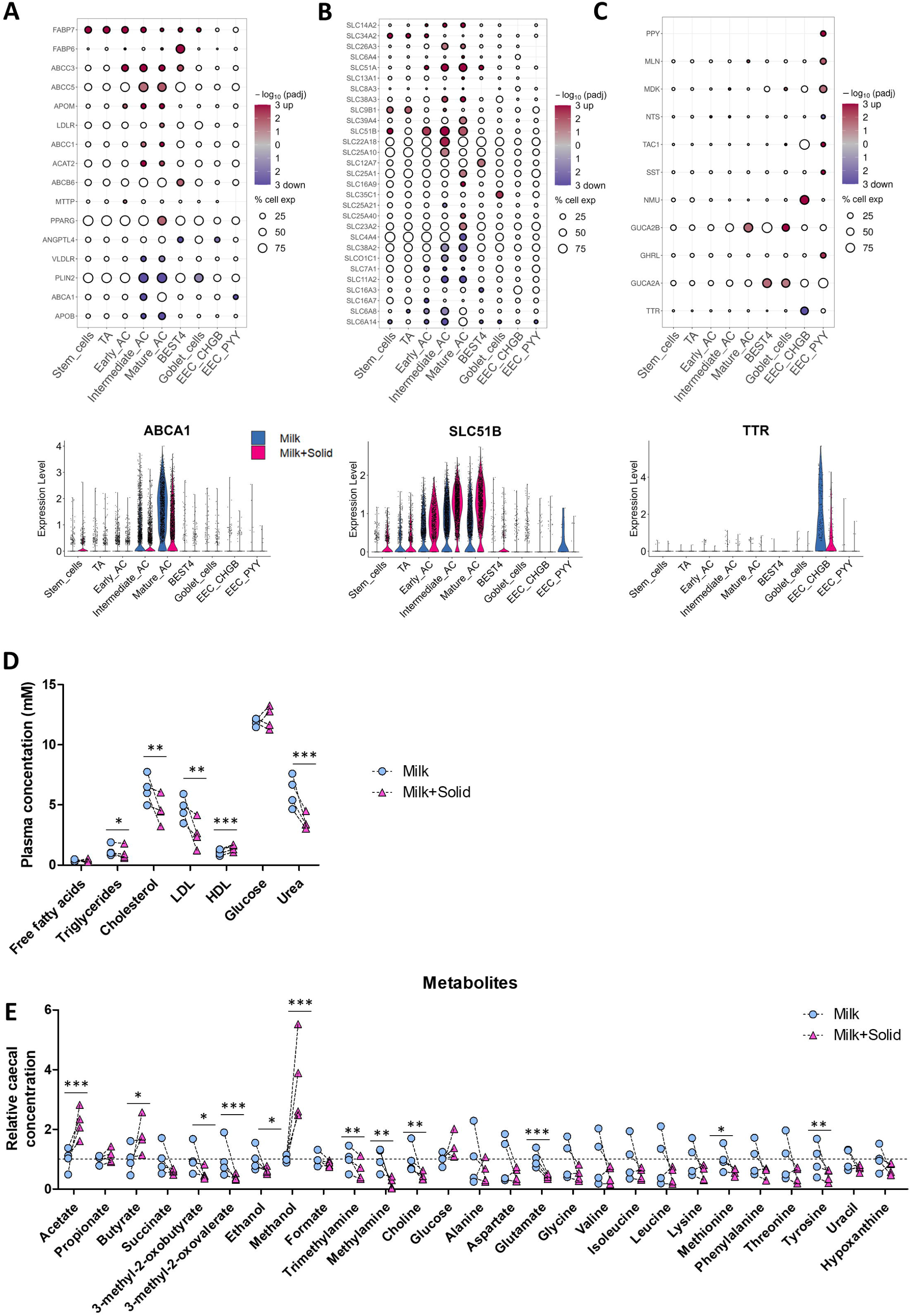
Solid food-induced modifications of the expression of genes involved in epithelial nutrient handling is associated with changes in concentrations of caecal and plasma metabolites. (A-C) Top: Selected differentially expressed genes (DEG) significance (log10(adjusted p-value) values, color) and fold change sign (red for over expressed and blue for under expressed genes in the “Milk+Solid” group versus the “Milk” group) involved in (A) lipid metabolism, (B) epithelial transport and (C) hormone secretion. The size corresponds to the percentage of cells expressing the gene in the cell type. Bottom: Expression level of a selected DEG by cell (dot), per cell type and group. (D) Plasmatic concentrations of metabolites. Points represent individual values per rabbit and dotted lines link littermates. *: p-value < 0.05, **: p-value < 0.01, ***: p-value < 0.001. HDL: high-density lipoprotein; LDL: low-density lipoprotein. (E) Relative caecal concentrations of metabolites detected by nuclear magnetic resonance-based metabolomics. Points represent individual values per rabbit and dotted lines link littermates. *: adjusted p-value <0.05, **: adjusted p-value <0.01, ***: adjusted p-value <0.001. AC: absorptive cells, EEC: enteroendocrine cells, TA: transit amplifying cells

In addition, the introduction of solid food altered the expression of numerous genes coding for SLC carriers (Figure 7B). Ingestion of solid food strongly upregulated the expression of genes coding for the urea transporter *SLC14A2* and several ion transporters (e.g., *SLC11A2, SLC22A18, SLC26A3, SLC39A4)* in absorptive cells (Figure 7B). The upregulation of the urea transporter coincided with a significant decrease in plasma urea concentration following the introduction of solid food (Figure 7D). Solid food ingestion also specifically upregulated the ion transporter *SCL12A7* in BEST4^+^ cells and the fucose transporter *SLC35C1* in goblet cells. The introduction of solid food altered the expression of monocarboxylic transporters with an upregulation of *SLC16A9* in mature absorptive cells and a downregulation of *SLC16A3* in BEST4^+^ cells and of *SLC16A7* in early absorptive cells (Figure 7B). This result can be linked to the increased concentrations of the bacterial short chain fatty acids acetate and butyrate in the caecum content after the introduction of solid food (Figure 7E). Similarly, the lower concentration of amino acids (glutamate, methionine, and tyrosine) following solid food introduction could be linked to the upregulation (*SLC38A3)* and downregulation (*SLC38A2, SLC6A14, SLC7A1*) of amino acids transporters in absorptive cells (Figures 7B and 7E).

The solid food-induced alterations of nutrient handling were associated with modifications of gene expression in EEC (Figure 7C). In EEC PYY^+^ cells, solid food introduction upregulated the expression of genes coding for the hormones *GHRL*, *MDK, MLN, SST, PPY,* and *TAC1*, while only *NTS* was downregulated. In *EEC CHGB^+^*, solid food ingestion increased the expression of the hormone coding gene *NMU*, while *TTR* was downregulated (Figure 7C). These observations are reflected by the EEC-specific enrichment of DEG involved in the biological pathway “digestion” (Figure 4K). In addition, genes coding for the hormones involved in the guanylate cyclase C signaling (*GUGA2A* and *GUCA2B*) were upregulated in goblet cells, mature absorptive cells and BEST4^+^ cells (Figure 7C). Overall, our results show that solid food ingestion enhances epithelial differentiation and remodels the sensing, transport and metabolism of nutrients by epithelial cells.

## Discussion

Our study provides the first single-cell transcriptomic atlas of the rabbit intestinal epithelium. This dataset expands the characterization of the cellular diversity of the intestinal epithelium in mammals and constitutes an important resource for the use of rabbits as a model in gastrointestinal research. Our results notably highlighted the diversity of absorptive epithelial cells in the caecum. Indeed, we observed a functional specialization of absorptive cell subsets along the caecal crypt axis that mirrored previous findings in the small intestine villi (Moor et al. 2018; Beumer et al. 2022). For instance, our results showing that middle-crypt absorptive cells specifically express genes coding for antimicrobial peptides is consistent with the observation made in bottom villus enterocytes (Moor et al. 2018; Beumer et al. 2022). Another important finding of our study is the high homology between rabbit and human BEST4^+^ cells, which suggests that the rabbit could be an appropriate animal model to study the role of this subset of mature absorptive cells that are absent in mice (Malonga et al. 2024). Additionally, the two subsets of EEC that we identified in the rabbit caecum (EEC PYY^+^, corresponding to L-cells expressing *GCG* and *PYY*; EEC CHGB^+^, corresponding to enterochromaffin cells expressing *TPH1*) were highly similar to human EEC, notably because EEC from both species express the hormones *MLN, MDK*, and *PPY* which are not expressed by mouse EEC (Parikh et al. 2019; Beumer et al. 2020). The absence of M cells in our caecal epithelium dataset was expected as these cell types are known to be present only in the small intestine (Burclaff et al. 2022; Schumacher 2023). The lack of Paneth and tuft cells in our single-cell survey could be explained by their scarcity, which reduces their probability of capture by droplets in the microfluidic system (Silverman et al. 2024). Indeed, our electron microscopy observations revealed a rare population of Paneth-like cells in rabbit caecal epithelial crypts, confirming previous reports (Cui et al. 2023).

The maturation of the intestinal epithelium at the suckling-to-weaning transition is considered to be largely driven by genetically wired factors while the contribution of nutritional and microbial signals remains debated (Muncan et al. 2011; Navis et al. 2019; Beaumont et al. 2022; Díez-Sánchez et al. 2024). Our results clearly demonstrate that the introduction of solid food is sufficient to induce major transcriptome modifications in every intestinal epithelial cell type, independently of age-related factors. Importantly, we observed this strong epithelial response to solid food despite the level of milk intake remaining unchanged, which indicates that the loss of milk-derived factors is not mandatory to induce epithelial maturation. Several previous studies described the transcriptomic changes occurring in the intestinal epithelium at the suckling-to-weaning transition (Rakoff-Nahoum et al. 2015; Pan et al. 2018) but, to our knowledge, our study is the first to reveal in which cell types these modifications take place. For instance, we newly demonstrated that the well characterized up-regulation of the immunoglobulin transporter *PIGR* at the onset of solid food ingestion (Jenkins et al. 2003; Rakoff-Nahoum et al. 2015; Beaumont et al. 2020) mainly occurs in epithelial cells localized at the crypt base (stem cells, TA cells, and early absorptive cells). This zonated expression of *PIGR* in crypt base cells that we confirmed by RNA *in situ* hybridization could be explained by the proximity with the underlying IgA secreting plasma cells. IgA secretion by crypt base cells could contribute to protect the stem cell niche from microorganisms. In line with our findings, the compartmentalization of *PIGR* expression in the intestinal epithelium was recently demonstrated to be driven by BMP signaling, which increases from the crypt base to the top (Beumer et al. 2022). Interestingly, we also observed an upregulation of *PIGR* expression in goblet cells after the ingestion of solid food, which suggests that transepithelial transport of IgA could be an unexplored function of mucus secreting cells. The presence of IgA in rabbit caecal goblet cells, as observed previously in the intestine of birds (Bar-Shira et al. 2014), could be explained by the binding of IgA to mucins or their transport through goblet cell-mediated passage (Knoop and Newberry 2018). Future research is required to explore the potential contribution of goblet cells to IgA transport across the intestinal epithelium.

Although our results indicate that most transcriptome changes induced by solid food ingestion are cell type-specific, we also found a few genes similarly regulated in most epithelial cell types. A striking example is the pan-epithelial upregulation of *ALDH1A1*, which is involved in epithelial processing of dietary vitamin A into retinoic acid (Bang 2023). Epithelial retinoic acid metabolism was previously shown to be upregulated at the weaning transition in the mouse intestine (Layunta et al. 2021) and is known to be influenced by the gut microbiota (Bang 2023), notably through the bacterial metabolite butyrate, which is able to induce *ALDH1A1* expression in epithelial cells (Schilderink et al. 2016). Accordingly, we observed that the solid food induced upregulation of *ALDH1A1* coincided with an increased concentration of butyrate and a higher abundance of the butyrate-producing family *Lachnospiraceae* (Vacca et al. 2020), which is probably driven by the introduction of plant-based complex carbohydrates. Given the role of retinoic acid in tuning intestinal immune responses (Hall et al. 2011), our results suggest that epithelial regulation of vitamin A metabolism at the onset of solid food ingestion may contribute to the “weaning reaction”, which corresponds to a transient remodeling of mucosal immunity essential to program mucosal health (Al Nabhani et al. 2019). In our study focusing on the epithelial layer, we found that a prominent feature of this “weaning reaction” was the strong upregulation of ISG, which was previously shown to be a microbiota-dependent process (Rakoff-Nahoum et al. 2015). Our study newly shows that this solid food induced upregulation of ISG is mostly restricted to crypt-top mature absorptive cells and to BEST4^+^ cells. This observation is in line with previous studies in mice showing that microbial colonization induced the upregulation of ISG specifically in subsets of mature absorptive cells localized at the tip of epithelial villi (Sommer et al. 2015; Van Winkle et al. 2022). In contrast with the upregulation of ISG, we found that solid food ingestion reduced the expression of numerous cytokines and antimicrobial peptides in a cell type specific manner, indicating an overall remodeling of epithelial defense systems. For instance, our data revealed that BEST4^+^ cells are the main producers of the immunomodulating *IL33* alarmin (Schiering et al. 2014), which is downregulated after ingestion of solid food. We also confirmed the goblet cell-specific expression of the recently discovered antimicrobial peptide *WFDC2* (Parikh et al. 2019) and we newly report its downregulation after the introduction of solid food.

Our results showing that solid food ingestion alters the gene expression of membrane mucins (*MUC1, MUC13, MUC12*), specifically in absorptive and BEST4^+^ cells highlight the cell types involved in the establishment of the glycocalyx, which was previously shown in mice to be part of an adaptation of the epithelial defense repertoire during weaning (Layunta et al. 2021). In addition, we found that several glycosyltransferases involved in the post-translational modification of mucins were regulated by the introduction of solid food predominantly in stem cells and proliferating cells located at the crypt-base. This effect may be driven by the changes in the gut microbiota induced by solid food ingestion since a previous study showed that microbial colonization of the mouse intestine similarly changed glycosylation in stem and transit-amplifying cells (Tsang et al. 2022). The mature absorptive cell-specific upregulation of the fructosyltransferase *FUT2* induced by the ingestion of solid food could also be driven by microbial signals but also by changes in glucocorticoid levels at the weaning transition (Nanthakumar et al. 2013; Arike et al. 2017). The solid food induced upregulation of mucus components secreted by goblet cells (*ZG16, TFF2*) could contribute to protect the intestinal epithelium from microorganisms expanding in the gut at the weaning transition.

The remodeling of epithelial defense systems induced by the introduction of solid food coincided with a shift of the transcriptome of caecal epithelial cells characterized by a reduced expression of small intestine-specific genes (*ANPEP, APOB*) and a higher expression of large intestine-specific genes (*AQP8, CA1, SLC26A3*) (Burclaff et al. 2022). The changes in the gut microbiota induced by solid food ingestion could contribute to the acquisition of these large intestine-specific functions since microbial colonization of the rat intestine was previously shown to induce similar effects (Tomas et al. 2015). The regional specialization of epithelial cells upon solid food introduction was associated with a strong shift in amino acid and lipid metabolism. Indeed, solid food introduction altered the absorptive cell expression of transporters of amino acids whose concentration was reduced in the lumen. Increased utilization of milk-derived amino acids by the microbiota for bacterial growth could lower their availability (Portune et al. 2016), which could explain the lower urea concentration in the plasma and the increased expression of its transporter (*SLC14A2*) in absorptive cells after the introduction of solid food. The effects of solid food ingestion on the expression of lipid handling genes in absorptive cells could also be driven by changes in the gut microbiota, which has previously been shown to regulate lipid homeostasis in the intestinal epithelium during the weaning transition in mice (Rakoff-Nahoum et al. 2015). Indeed, metabolites produced by the gut microbiota in early life regulate lipid metabolism in epithelial cells (Araújo et al. 2020; Shelton et al. 2023). In contrast, changes in dietary lipid supply can be ruled out, as lipids are mainly derived from maternal milk (Gidenne and Fortun-Lamothe 2002), the amount of which was not reduced after the introduction of solid food. Changes in the gut microbiota triggered by solid food ingestion could also contribute to the upregulation of basolateral bile acid exporters (OSTα/β coded by *SCL51A/B* genes) in absorptive cells and to the slight reduction of the plasmatic concentration of cholesterol, the precursor of bile acids (Rakoff-Nahoum et al. 2015; Le Roy et al. 2019; Collins et al. 2023). In turn, solid food-induced modification of bile acid metabolism could contribute to the maturation of the microbiota, as demonstrated in mice at the suckling-to-weaning transition (van Best et al. 2020). Interestingly, we found that the cytosolic bile acid binding protein (*FABP6*) was specifically expressed and upregulated in BEST4^+^ cells after solid food ingestion, which suggests an uncovered role for these cells in the enterohepatic circulation (Dawson and Karpen 2015).

Our study is focused on epithelial cells, whereas major changes are known to occur in intestinal immune cells during the weaning transition (Al Nabhani et al. 2019). Our initial scRNA-seq dataset included some intraepithelial lymphocytes but their numbers were insufficient to perform reliable analyses. Future studies analyzing the single-cell transcriptome of *lamina propria* immune cells in our suckling rabbit model ingesting or not solid food are needed to expand our understanding of the gut barrier maturation during the weaning transition. Indeed, the transcriptome changes that we observed in intestinal epithelial cells after solid food ingestion suggest alterations in the crosstalk with immune cells, particularly in relation to interferon and cytokine signaling. Another limitation of our study is related to its restriction to epithelial cells isolated from the cecum, which we chose because this gut region harbors a dense microbial population that is highly responsive to dietary changes at the suckling-to-weaning transition (Beaumont et al. 2020). Additional studies examining single-cell transcriptome changes induced by solid food ingestion in other regions of the gut, such as the jejunum, may also be relevant to explore metabolic and immune modulations. Furthermore, previous studies have shown that microbiota changes are directly involved in epithelial bulk transcriptome modifications at the weaning transition (Rakoff-Nahoum et al. 2015; Pan et al. 2018), while our study performed at the single-cell level did not evaluate this causal role. The recent development of intestinal organoids that recapitulate the cellular diversity of the epithelium *in vitro*, including in rabbits (Mussard et al. 2020), will be useful in future studies aiming at evaluating the cell type specific transcriptome changes induced by gut bacteria or metabolites modified at the weaning transition.

## Conclusion

In conclusion, our study provides the first single-cell transcriptomic atlas of the rabbit intestinal epithelium and significantly expands the understanding of cellular diversity in the mammalian intestine. We highlighted the homology between rabbit and human intestinal epithelial cells, such as BEST4^+^ cells, supporting the suitability of the rabbit as a model for gastrointestinal research. In addition, we uncovered cell type specific transcriptome modifications driven by solid food ingestion at the suckling-to-weaning transition, highlighting changes in epithelial defense mechanisms and metabolic processes. These findings contribute to a broader understanding of the postnatal maturation of the gut barrier in mammals.

## Acknowledgments

This work was supported by a grant from the French National Research Agency: ANR-JCJC MetaboWean (ANR-21-CE20-0048). Tania Malonga was supported by grants from Toulouse-INP, INRAE and Oak Ridge Institute for Science and Education (ORISE). The authors thank Isabelle Fourquaux (CMEAB, Toulouse, France) for assistance with electron microscopy analyses. We acknowledge the I2MC cytometry and cell sorting facility (Genotoul-TRI), member of the national infrastructure France-BioImaging supported by the French National Research Agency (ANR-10-INBS-04).

## Disclosures

The authors declare no conflict of interest.

## Author contributions

CK, CLL, NV and MB designed research;

TM, CK, EL, AP, EJ, CL, MD, IP, ER, AI and MB conducted research;

TM, CC, NV and MB analyzed data;

TM, NV and MB wrote the initial draft.

All authors have read and approved the final manuscript.

## Notes

### Competing Interest Statement

The authors have declared no competing interest.

### Summary of Updates

Affiliations of the authors were corrected

