## Supplementary figures and images for "A single-cell atlas of transcriptome changes in the intestinal epithelium at the suckling-to-weaning transition"

### Figure S1

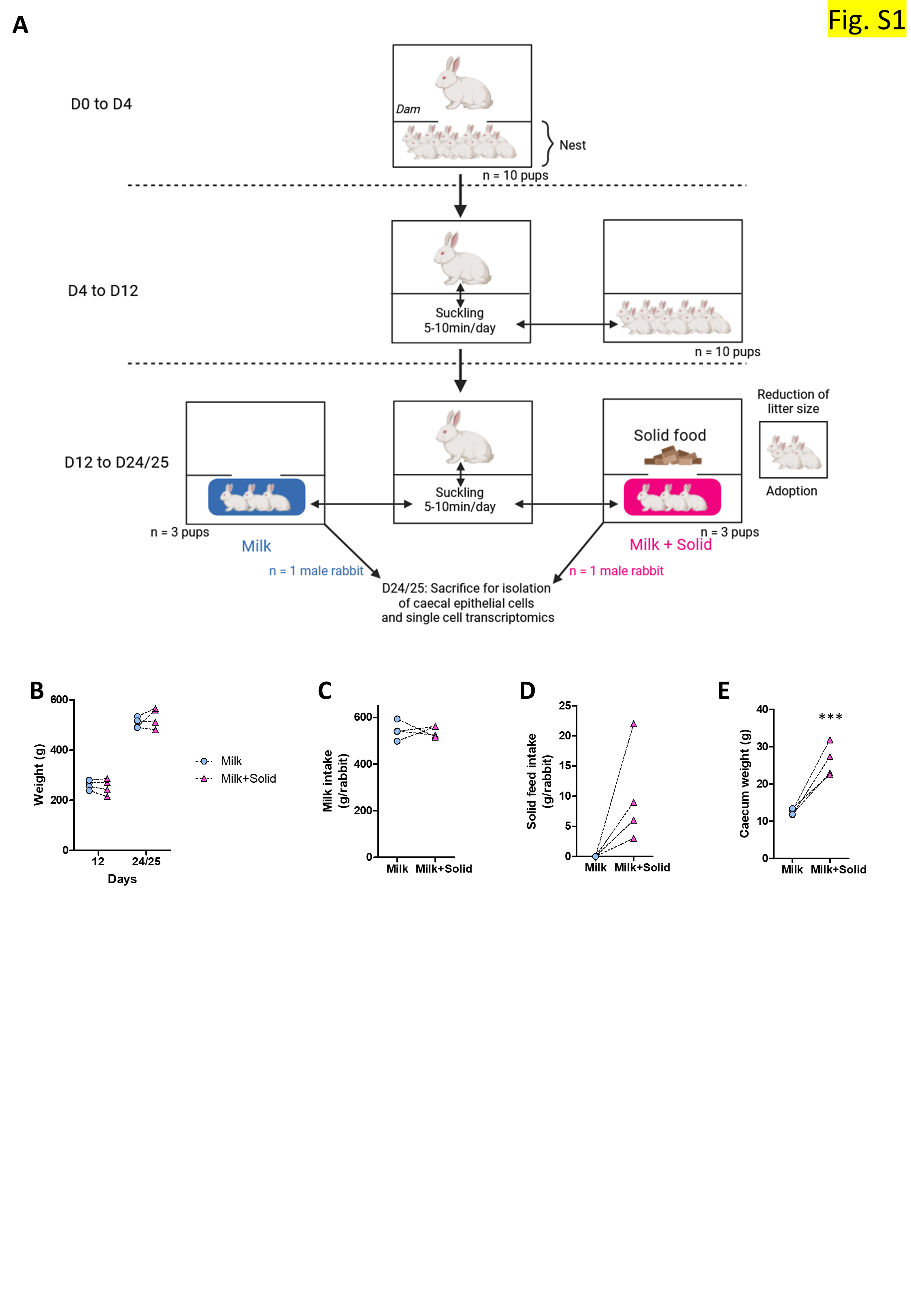

### Figure S2

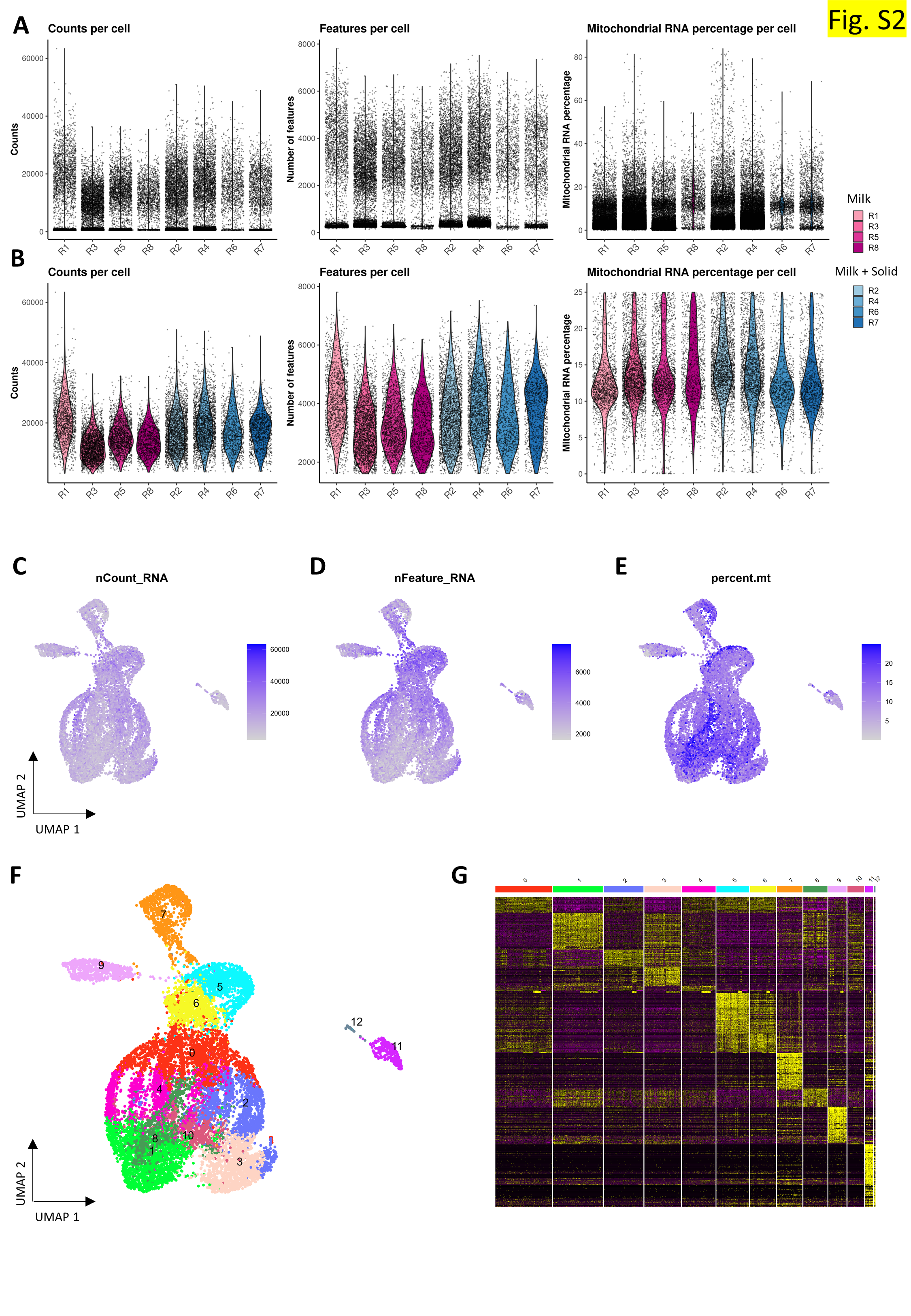

### Figure S3

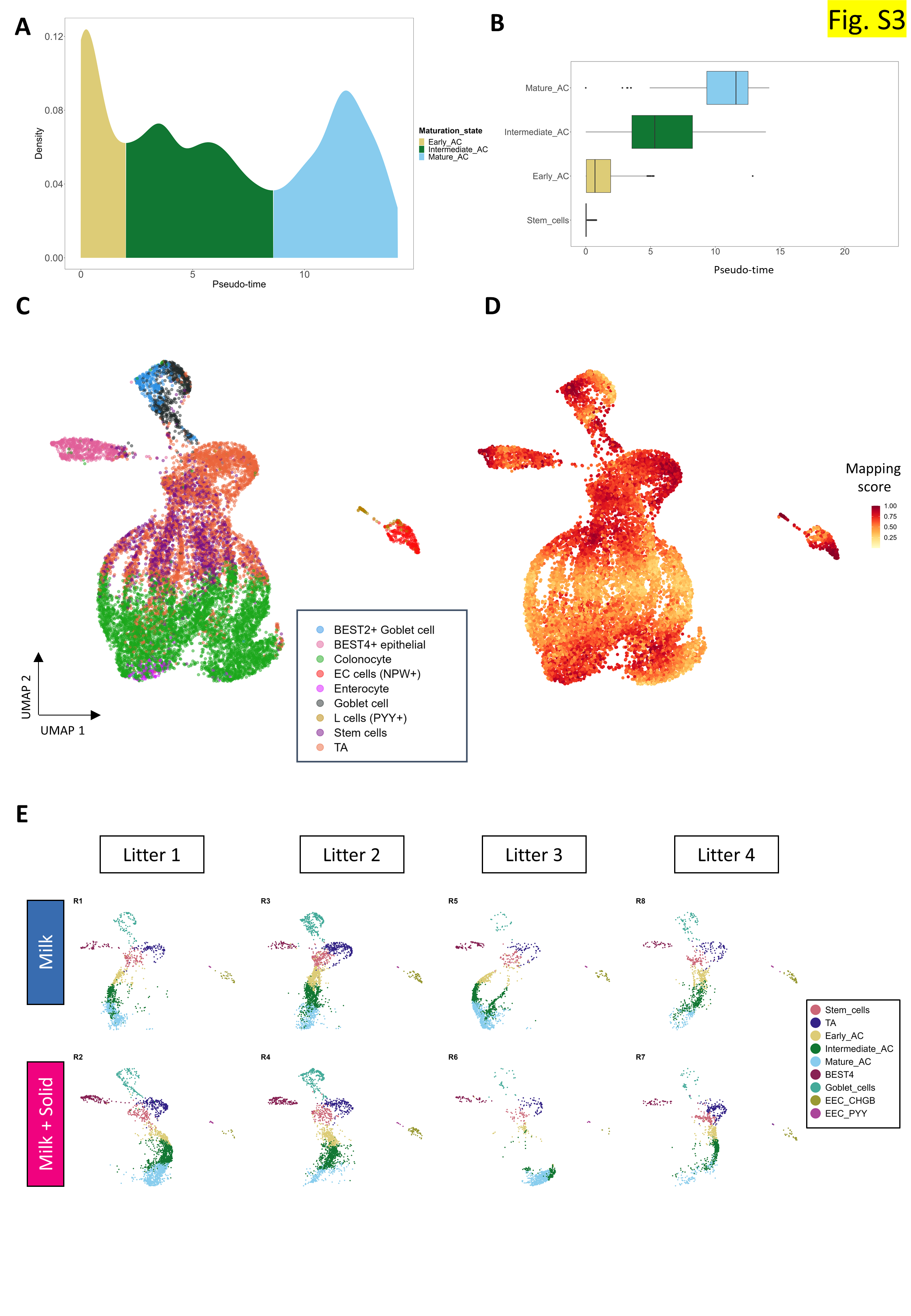

### Figure S4

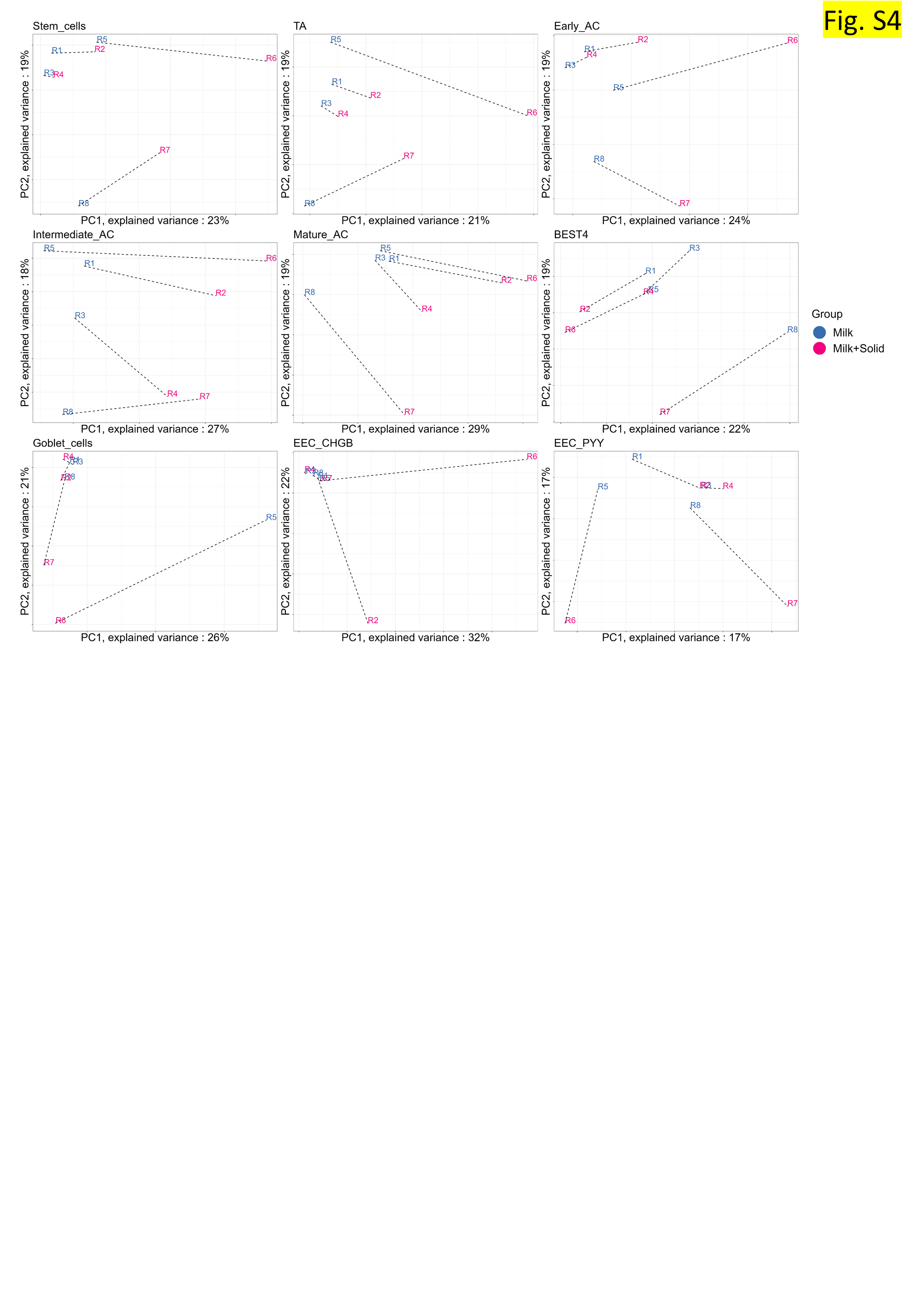
